# Micellangelo: a Generative Model of Cell-Topography Interactions

**DOI:** 10.1101/2025.11.24.690184

**Authors:** Nikita Konshin, Koen Minartz, Jan de Boer, Vlado Menkovski

## Abstract

Understanding how materials influence cell behavior is central to biomaterial engineering, yet experimental approaches are limited by throughput and complexity. Here, we present Micellangelo, a generative AI model that simulates high-resolution fluorescence images of cells cultured on micro-topographical surfaces. By conditioning on surface topographies, Micellangelo generates realistic cell morphologies, enabling in silico exploration of cell–material interactions. We fabricated a dataset of human dermal fibroblasts stained for DNA, F-actin, and the mechanosensitive transcription factor YAP across ten distinct topographies. Micellangelo was trained using the flow matching framework, and demonstrated strong visual and quantitative alignment with real microscopy data in terms of morphology, intensity patterns, and mechanotransductive features. Beyond image generation, we demonstrate two proof-of-concept in silico experiments: tuning the strength of topographical conditioning and perturbing subcellular structures to assess phenotypic consequences. The model captures both topography-induced morphological trends and structure–function relationships such as actin–YAP coupling. These results suggest that Micellangelo functions as a digital twin of experimental cell imaging, taking a step towards a scalable platform for hypothesis generation, virtual screening, and design of biomaterial interfaces. This work bridges the fields of biomaterials and generative modeling, and introduces a generalizable framework for conducting biologically meaningful in silico experiments.

## 1. Introduction

Understanding how materials influence cells is crucial for both fundamental cell biology and for the development of effective biomedical applications, such as medical implants and drug delivery systems.^[1–9]^ Cell-material interactions involve a broad range of material properties, including surface chemistry, stiffness and topography, all of which can influence cellular physiology and fate. To explore the biomaterial design space, biomaterial engineering has undergone a transformative evolution from early, small-scale experimentation to expansive high-throughput screening (HTS) platforms.^[10–16]^ Using HTS technology, a large variety of material designs can be scanned in parallel. Still, even HTS remains constrained by finite laboratory capacity and resources, which stands in stark contrast to the virtually infinite and combinatorial design space of biomaterials. This has led towards the development of computational models, allowing researchers to scan even larger ranges of material designs in silico before moving to laboratory experiments. The ultimate use case of such computational approaches is in silico experimentation, where cell responses to material cues are simulated directly on the computer.^[17,18]^ This empowers researchers to prototype surface designs, test hypothetical scenarios, and perform experiments that are costly or technically infeasible in the lab. We can see similar approaches in other fields such as drug discovery, ^[19]^ where virtual screening and perturbation modeling are widely used thanks to standardized datasets and well-annotated outcomes.

However, unlike computational models in physics and chemistry that can simulate systems in their entirety based on first principles,^[20]^ it remains challenging to approximate all aspects of cell behavior in an integrated simulation using computational models based on physics-like laws. Instead, computational approaches in cell biology often rely on statistical models that predict a single quantity of interest, for example cell proliferation, based on prior experimental data.^[16,21–24]^ Notably, such models reduce the biological experiment to a single output value. This reductionist approach precludes interpretation of the outcome by cell biologists, which relies on visually inspecting the complete experiment. Consequently, although such predictive models have proven successful, true in silico experimentation requires a fundamentally different approach that is capable of modeling not just experimental results, but also the experimental process that led to them. In the context of imaging-based biomaterials experiments,^[25,26]^ this amounts to simulating fluorescence microscopy images conditioned on the material properties of interest.

Over the past decade, developments in deep generative models have unlocked the capability to generate new data from complex, high-dimensional distribution.^[27–33]^ Initially focused on data such as images and text, the generative modeling paradigm is well-suited for in silico surrogate experiments, as it aligns naturally with the goal of generating full, high-dimensional experimental outcomes, such as microscopy images, conditioned on experimental parameters. Accordingly, this approach has started to gain traction in the cell biology community over the last years. For example, Generative Adversarial Networks and diffusion-based models have been proposed for expressing how chemical perturbations affect cell phenotype. (Bourou, Mahanta, et al., 2025a; Bourou, Segade, et al., 2025a; Bourou Anisand Boyer, 2024a; Lamiable et al., 2023a; Zhang et al., 2025a)

However, applying generative models to biomaterials comes with unique challenges, as modeling experimental outcomes requires characterizing the effect of a diverse range of material cues, among which chemical, mechanical and geometric properties.^[39]^ In this paper, we present Micellangelo, a generative model capable of synthesizing high-resolution cell images based on underlying surface topographies, advancing the field toward this goal. Surface topography stands out as particularly amenable to modeling: it is parameterizable and can be visually described, making it a practical entry point for image-based generative modeling in the field of biomaterials. Moreover, the TopoChip platform, a high-throughput system that uses microfabricated polystyrene surfaces with engineered topographies, facilitates HTS to gather large amounts of training data. Each topography exerts unique biophysical influences onto cells, modulating properties such as spreading, polarity, cytoskeletal organization, and nuclear architecture.^[23,40]^ We collected a dataset of normal human dermal fibroblasts (referred to as fibroblast) stained for DNA, F-actin and the mechanosensitive transcription factor YAP on ten distinct topographies. By integrating spatially resolved material cues with generative modeling, Micellangelo offers a scalable framework for computationally reproducing cell–material interactions. Whereas prior approaches in this domain have focused on statistical associations or predictive classifiers, Micellangelo can generate full cellular morphologies informed by the biomaterial design. This opens the door to new research directions in virtual material screening, design optimization, and biomaterial interfaces based on large-scale HTS data, where one can iterate on material design and conduct biological experiments entirely in the digital domain.

## 2. Results

### 2.1. Database of single cell images on topographies

To construct a high-quality dataset of cellular response to surface architecture, we fabricated and screened a curated library of polystyrene substrates featuring ten distinct micro-topographies. These designs were selected from a larger set of surfaces previously shown to cause a diverse array of cell morphologies and phenotype.^[41]^ Each topography was embossed into a 13 mm polystyrene diameter insert, allowing full-surface imaging at high resolution. The experimental workflow **(Supplementary Figure S1)** encompassed topography fabrication via hot embossing, cell seeding, multiplexed immunofluorescence staining, widefield microscopy, and downstream cell segmentation and feature extraction. To verify the fidelity of surface patterning, each insert was imaged using brightfield microscopy **(Supplementary Figure S2)**. Surface codes, such as 1130, 1330, 1710, and 2113, refer to unique micro-architectures differing in shape complexity, feature spacing, and orientation. As observed before, our microfabrication method produced consistent structural integrity across all inserts at high spatial resolution, but was not able to reproduce the sub-micron surfaces architecture present in some designs. ^[42]^

Fibroblasts were cultured for 48 hours on ten distinct polystyrene inserts containing engineered micro-pillared topographies, followed by fixation and multiplexed staining for DNA (DAPI), F-actin (phalloidin), and the mechanosensitive transcription factor YAP. On flat control surfaces, fibroblasts displayed elongated morphologies with centrally located nuclei and prominent actin stress fibers. In contrast, cells cultured on topographically patterned inserts exhibited distinct morphological phenotypes dependent on interpillar spacing **(Figure 1)**.

**Figure 1.**
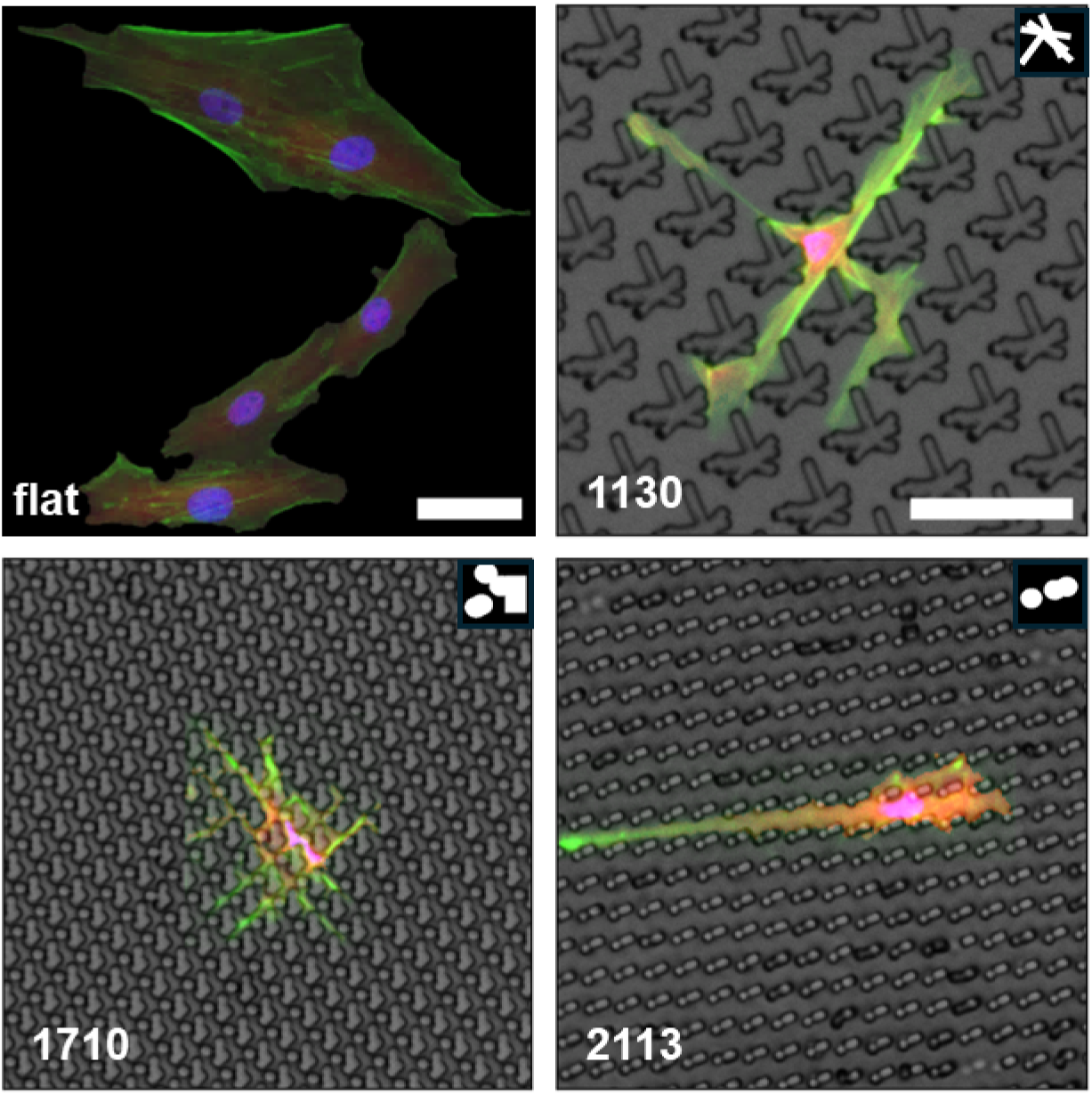
Fibroblast morphology in response to surface topography. Fibroblasts were cultured for 48 hours on flat or micro-pillared polystyrene inserts and stained for DNA (DAPI, blue), F-actin (phalloidin, green), and YAP (red). Flat surfaces supported typical elongated morphologies, whereas topographically patterned surfaces induced shape changes reflecting varying degrees of physical confinement. Scale bar: 50 μm. Numbers in the images (e.g., 1130, 1710, 2113) refer to the FeatureID values of specific topographies from the TopoChip library. The corresponding designs and metadata can be found at: https://github.com/cbite/TopoChip-analysis/blob/main/data/metadata/FeatureImages.zip On surface 1130, which featured widely spaced pillars (5–18 μm interpillar distance), cells were fully confined within grooves between the features, adopting elongated shapes with actin filaments aligned along the pillar axis. On surface 1710, where the pillars were more densely packed (1–6 μm spacing), fibroblasts exhibited partial confinement, often spreading over, rather than between, the pillars, and adopting more compact or branched morphologies. Surface 2113, a hybrid layout (2–6 μm spacing), induced asymmetric cell responses, with partial alignment to the topographical features on one side and unconfined spreading on the other. Notably, at inter-pillar distances below ∼2 μm, cells no longer consistently grew between features and instead bridged across pillars, indicating a confinement threshold that impairs pillar-guided spreading.

In addition to these global surface-level effects, local positioning within the same topography was found to exert significant influence on nuclear morphology. Overlays of DAPI-stained nuclei onto brightfield topographical maps revealed three qualitatively distinct regimes of confinement: *minimal confinement*, in which nuclei occupied open regions and retained a round, unperturbed morphology; *partial confinement*, where nuclei were compressed by adjacent pillars but not fully enclosed; and *full confinement*, marked by pronounced nuclear deformation due to tight spatial enclosure between microfeatures **(Figure 2)**. These confinement states were observed across multiple surface designs, including surfaces 1130 and 2088, and frequently co-existed within a single insert. Such local confinement effects underscore the importance of sub-surface spatial variation in modulating nuclear shape and, by extension, cytoskeletal organization and mechanotransductive responses.

**Figure 2.**
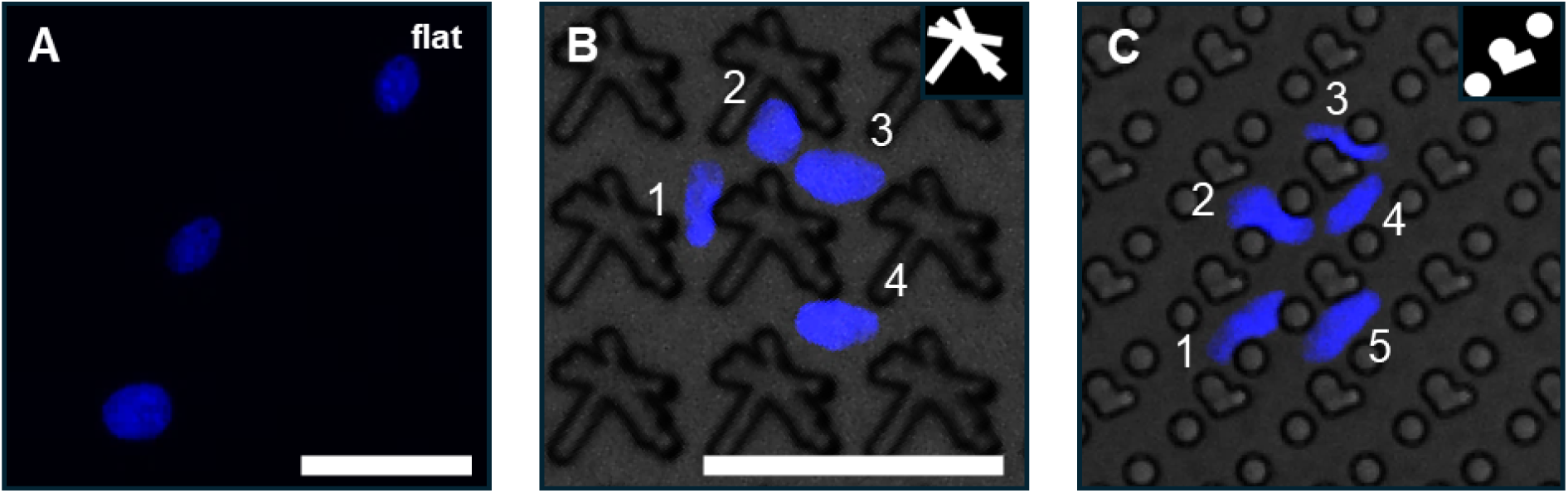
Effect of local topographical confinement on nuclear morphology. (A) Fluorescent DAPI signal (blue) of nuclei from cells cultured on flat polystyrene (PS), showing typical round morphology. (B) and (C) Brightfield images of two distinct surface inserts (B: surface 1130, C: surface 2088) overlaid with DAPI-stained nuclei from different positions. The overlays illustrate how cell position relative to surface topography alters nuclear shape. Three distinct types of nucleus–topography interactions are observed: Partial confinement, where nuclei are pressed by surrounding features but retain partially round morphology—examples: nucleus 1 and 2 in (B) and (C). Full confinement, where the nucleus is tightly enclosed between features, resulting in highly deformed shapes—example: nucleus 3 in (C). Minimal confinement, where the nucleus rests in open areas and retains a morphology similar to flat substrate—examples: nuclei 3 and 4 in (B), and 4 and 5 in (C). Scale bars: 50 μm.

Within a group of cells cultured on the same topography, we observed substantial heterogeneity in YAP localization across individual cells, a trend that was consistently present across all topographies. This variation suggests that while topography strongly modulates cell morphology and nuclear geometry, mechanosensitive signaling through YAP exhibits notable cell-to-cell variability. Together, these findings indicate that topography exerts a direct, heterogeneous influence on fibroblast morphology, nuclear architecture, and cytoskeletal organization, shaped not only by the global design of the substrate but also by the precise local spatial environment encountered by individual cells. The segmentation and quality control pipeline for single-cell extraction, including detailed illumination correction, threshold calibration, and classifier-based artifact removal, is described in **Supplementary Figures S3–S4**. To place each single cell in a consistent topographical frame, we defined a canonical coordinate system per insert. A clean 750×750 pixel brightfield snippet was chosen as the reference motif, and the brightfield channel of every single-cell crop was rigidly registered to this reference using phase cross-correlation. This registration yielded the cell’s offset within the repeating motif, enabling direct comparison of cells that experience the same local topographical context. Visual QC of the registered outputs confirmed correct relative alignment to the background pattern (see **Supplementary Methods**, **Standardized coordinate system**). In total, the curated dataset comprised 8,678 high-quality single-cell images collected from 13 surface topographies (10 unique designs), with a median of 682 ± 110 cells per surface (**Supplementary Table 3**).

### 2.2. In silico generation of cell images. Realistic single cell generation

After QC, we uploaded the images to a cluster equipped with an NVIDIA H100 GPU, trained Micellangelo as described in the Materials and Methods, and used the trained model to generate 8640 simulated cells. To generate cells, Micellangelo slowly transformed Gaussian noise to images of visually realistic cells **(Figure 3)**, following the flow matching framework upon which the model was built.^[32,43,44]^ We observed that Micellangelo first generates large-scale features of cell images, intuitively corresponding to rough outlines of the morphology, after which it focuses on more fine-grained details such as the texture of actin staining. After generation, we qualitatively investigated if the simulated cells visually resembled real cells that we observed in the lab **(Supplementary Figure S5)**. We noticed that the generated cells (Figure 4, bottom row) have similar visual characteristics as the real cells (top row), demonstrating qualitative alignment in terms of various characteristics relevant to fluorescent cell stain imaging, e.g. intensity, area, and shape. We also observed visual alignment in more fine-grained features, such as actin stress fibers, nucleus shape in relation to topographical confinement, and YAP intensity fluctuations **(Figure 4)**.

**Figure 3.**
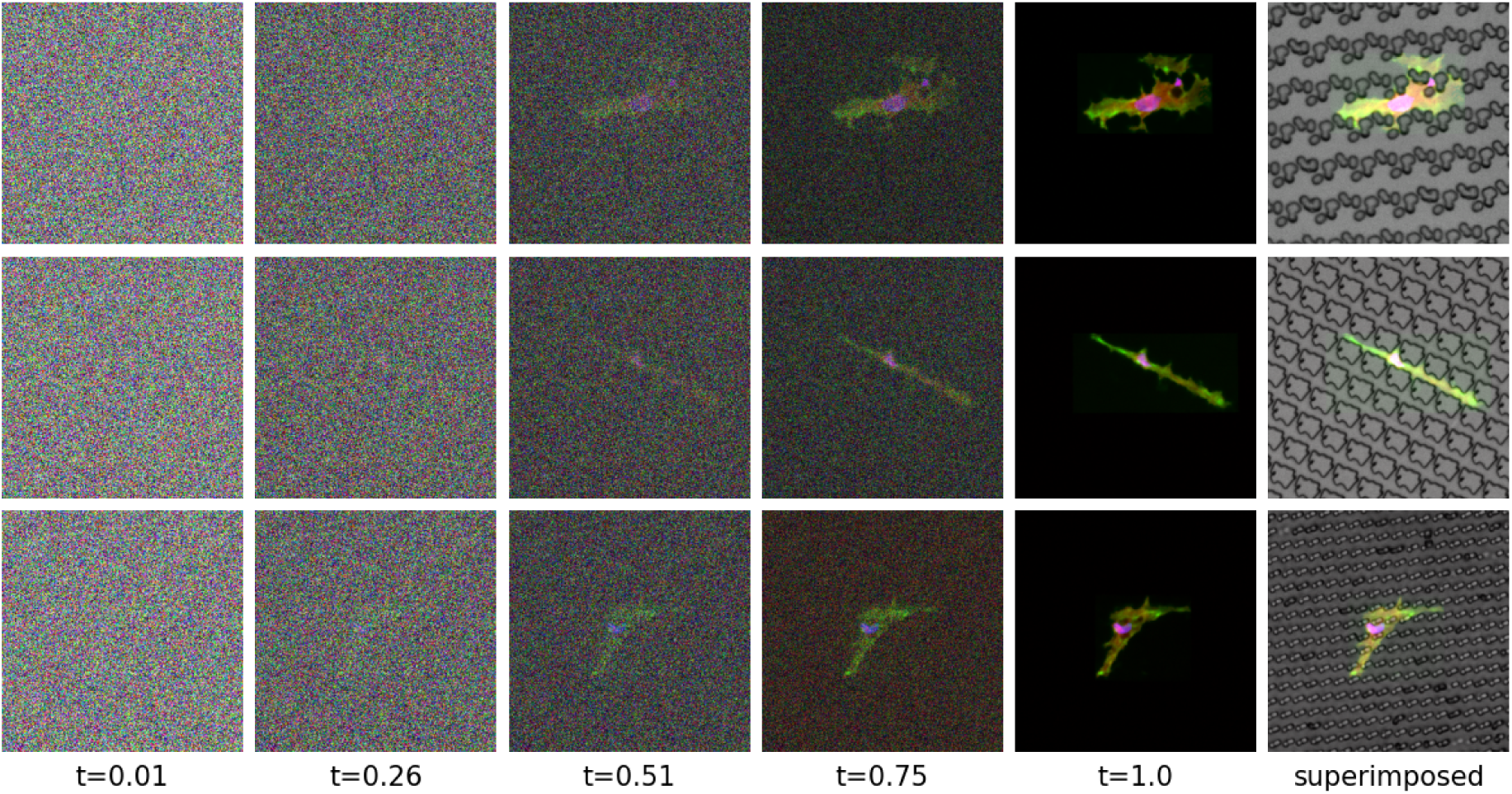
Generative transformation from Gaussian noise to realistic single-cell morphology using Micellangelo. Starting from Gaussian noise (t = 0), the image was gradually transformed into a biologically realistic cell image at t = 1 by integrating an ordinary differential equation defined by a vector field predicted by Micellangelo. This enabled the generation of high-resolution synthetic images conditioned on morphological and topographical context.

**Figure 4.**
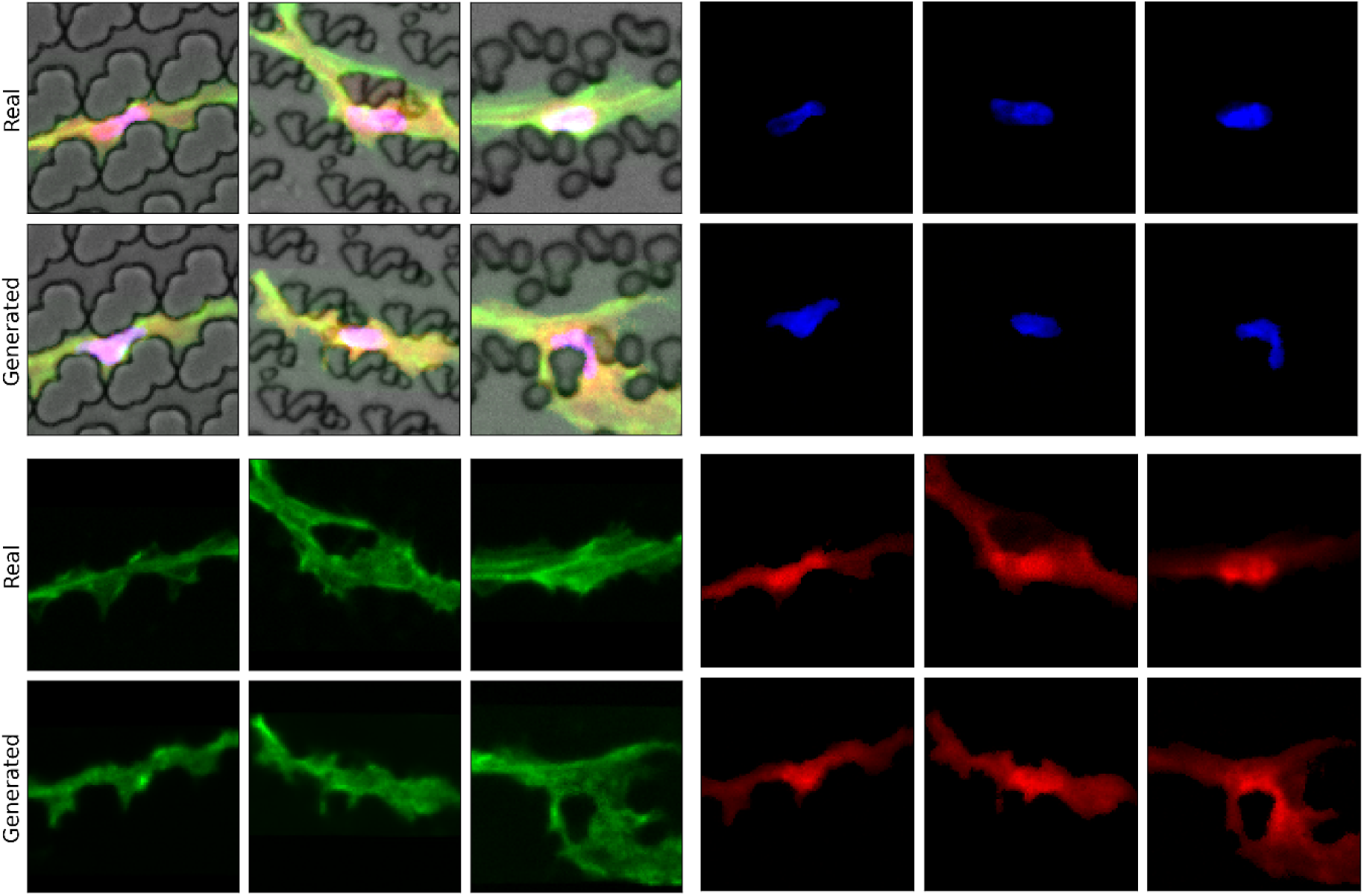
Comparison of individual fluorescence channels in real and Micellangelo-generated cells. For each pair, the top row shows cropped images of real fibroblasts, and the bottom row shows corresponding synthetic cells. The paired rows represent the composite image, DAPI (nuclei), phalloidin (actin), and YAP (mechanotransduction marker). The visual similarity across channels demonstrates the model’s ability to recapitulate fine-grained subcellular features such as intensity fluctuations.

### 2.3. Distribution alignment of cellular morphological features across surfaces

To quantitatively investigate whether the generated cells accurately resemble the observations made in the laboratory, both generated and real single-cell images were processed using a CellProfiler pipeline to extract relevant biological metrics; taken together, these define a morphological fingerprint of each cell. Since substantial morphological variability is expected even among cells cultured on the same topographical surface, we assessed Micellangelo’s accuracy by comparing the distributions of these metrics between real and simulated datasets. These distributions varied between given selected surfaces **(Figure 5)**, underscoring the role of topographical design in shaping cell behavior. For instance, DAPI roundness (form factor) was notably lower on surface 1710 compared to 1130, reflecting the more pronounced nuclear deformation due to strong physical confinement (see also Figure 3). Surface 1130 exhibited skewed distributions for both DAPI form factor and actin solidity, consistent with elongated, aligned cells squeezed between widely spaced pillars. Surface 1710 led to broader, more symmetric distributions, reflecting the partial confinement and diverse morphologies seen under higher pillar density. Meanwhile, surface 2113, with its hybrid architecture, led to asymmetric and highly concentrated cellular orientation. Importantly, across all three surfaces, the distributions generated by Micellangelo closely aligned with those of real cells, capturing not just average trends but also the surface-specific variation in shape, intensity, and organization.

**Figure 5.**
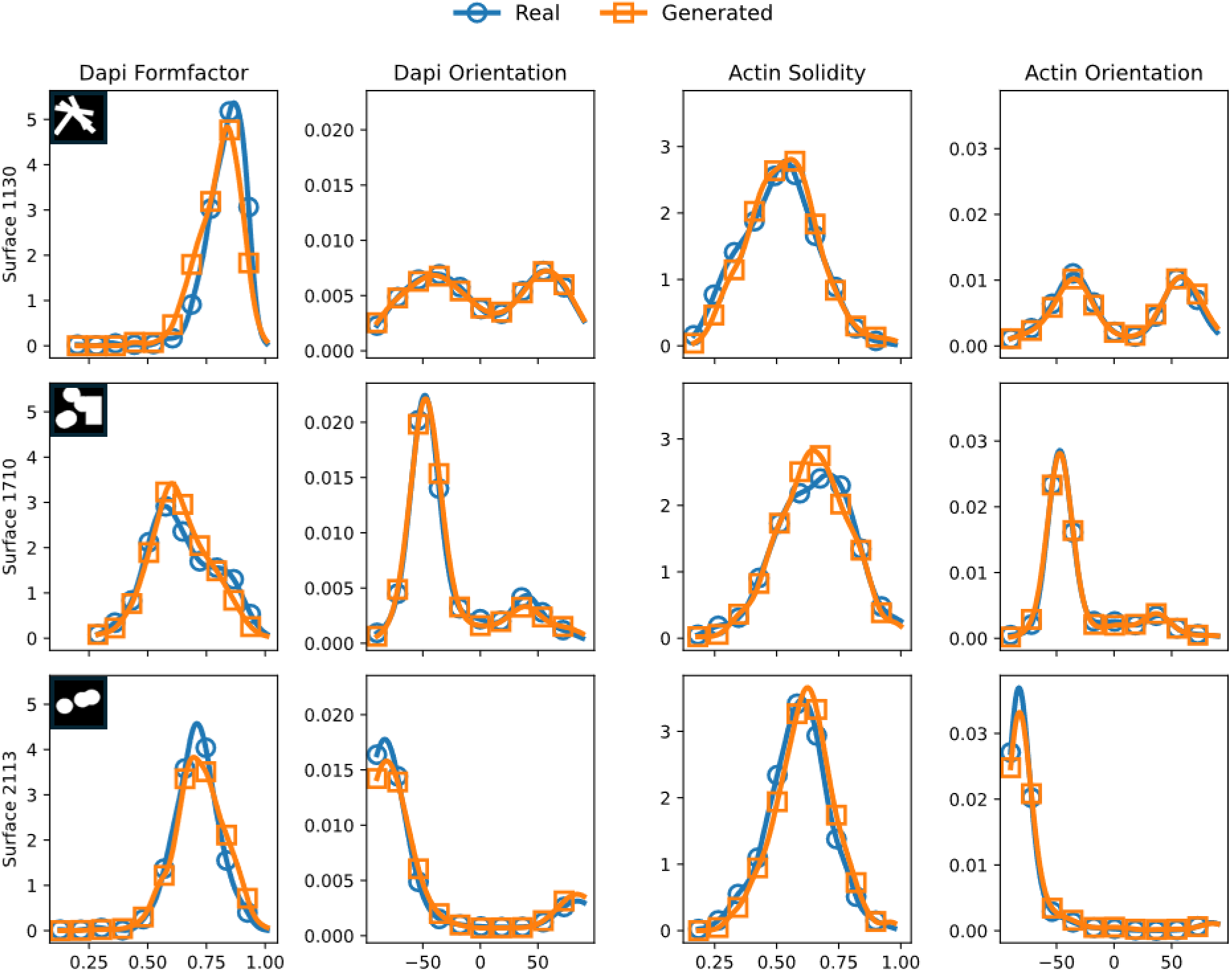
Morphological profiles of real and generated images. Probability distribution functions over various cellular metrics, calculated for surfaces 1130, 1710 and 2113 (top left corner of each row shows design schematics) for both real and generated cell image data. Distributions vary between surfaces, and generally the real and generated distributions overlap well for all surface-feature combinations. Y-axis shows probability density (frequency normalized) of cells in that metric bin — i.e., the fraction of cells per unit of the measured metric. *FormFactor* is defined as 4\**π*\*Area/Perimeter^2^. A value of 1 corresponds to a perfect circle; values closer to 0 indicate increasingly non-circular (irregular or elongated) object shapes. *Solidity* is defined as the ratio of object area to the area of its convex hull (ObjectArea/ConvexHullArea). A value of 1 means the object exactly fills its convex hull (i.e., has no indentations). Lower values indicate the object has concavities or is more irregular. *Orientation* is defined as the angle (in degrees, ranging from –90 to +90) between the x-axis and the major axis of the ellipse that has the same second-moments as the object. Thus it describes the tilt/rotation of the object relative to horizontal.

To further illustrate the variety over morphologies and the alignment of generated cells with real observations, we inspected the morphological fingerprint embedded in a two-dimensional space using tSNE **(Figure 6)**.^[45]^ Cells exhibited clear clustering by the surface they were cultured on, indicating distinct morphologies, and generated and real cells on the same surface were positioned close to each other, indicating morphological similarity.

**Figure 6.**
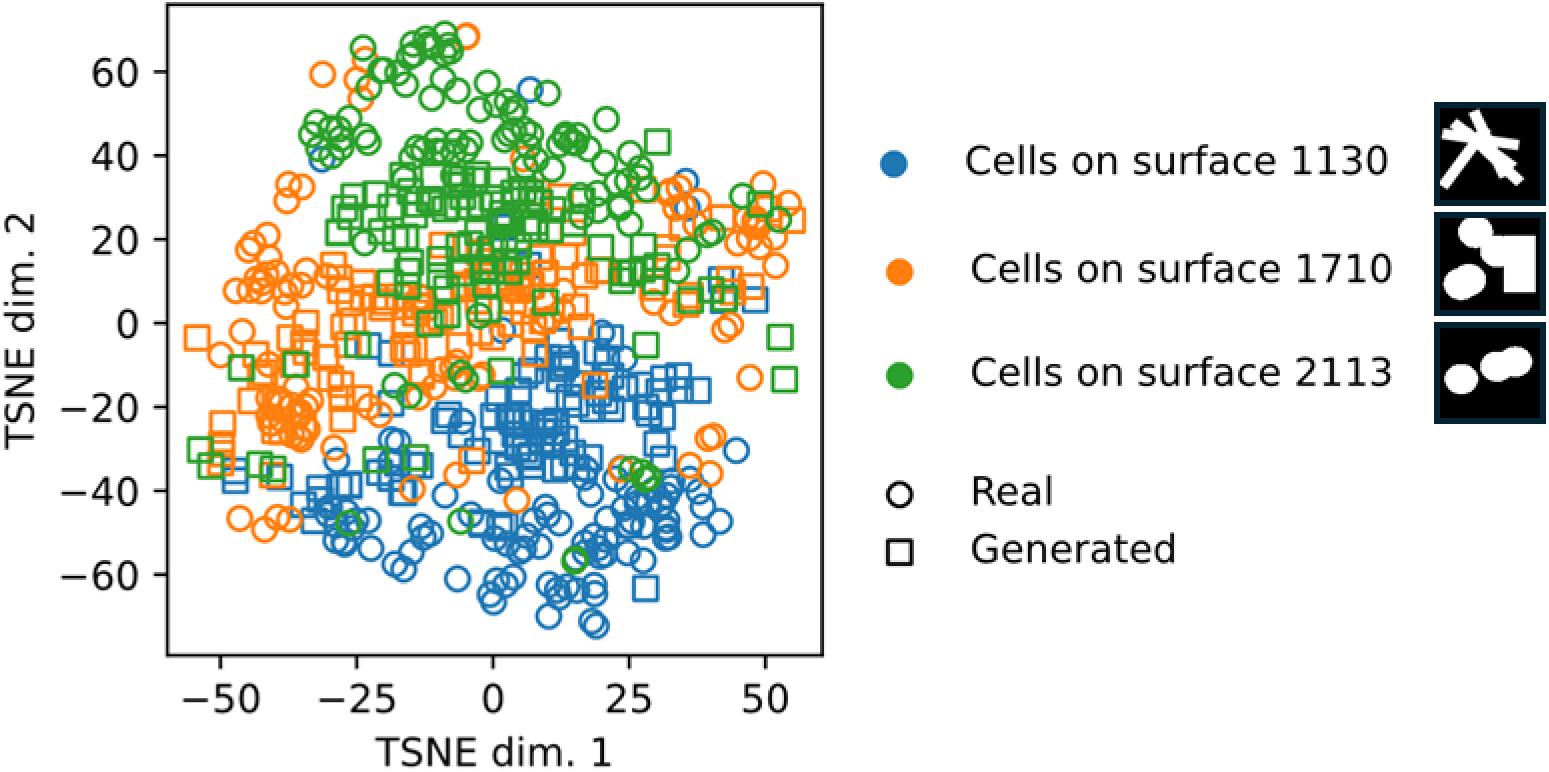
Similarity between real and generated cells. t-SNE embeddings of 376 morphological features describing cell shape and stain intensity (excluding location-related metrics). To reduce dimensionality, we first applied principal component analysis and retained the top 50 principal components, which were then used for t-SNE projection into two-dimensional space. Cells on different topographies form distinct clusters, reflecting the topography-dependent modulation of morphology. Real and generated cells occupy overlapping regions in embedding space, indicating that Micellangelo successfully reproduces surface-specific morphological patterns.

### 2.4. Modeling cellular structural-functional relationships

Having established alignment in the distribution of single morphological metrics, we investigated whether Micellangelo accurately captured relationships that are fundamental to cellular structural-functional organization. These correlations offer deeper insight into whether the model captured not just isolated traits, but also the biological interactions underlying cell behavior. We considered six representative feature correlations derived from both real and Micellangelo-generated cells **(Figure 7)**. These relationships include dependencies between nuclear shape (DAPI intensity and form factor), cytoskeletal morphology (actin area and intensity distribution), and mechanosensitive signaling (YAP localization). Overall, we found that Micellangelo reproduced key trends observed in real data. For example, in the top-left panel of Figure 10, a positive correlation was seen between DAPI eccentricity and upper quartile DAPI intensity, suggesting that more elongated nuclei tend to have slightly higher chromatin condensation, a pattern which emerged in both real and generated data and observed previously.^[46]^ Similarly, actin area correlated positively with DAPI area (top center panel), reflecting global scaling relationships in cell architecture, i.e. a cell that spreads will also flatten the nucleus thus leading to larger nuclear area. Importantly, the model also replicated biologically interpretable inverse correlations, such as the one between DAPI form factor and actin intensity heterogeneity (top right). This reflects the association of more pronounced actin bundles with less circular nuclei, in line with earlier literature on cytoskeletal–nuclear mechanical coupling.^[47–49]^ In the bottom row, three panels illustrate how actin area and shape are related to YAP intensity. We observed that median nuclear YAP intensity (bottom left) increased with actin area while cytoplasmic YAP (center) remained approximately constant, and that median nuclear YAP decreased as a function of actin form factor (right). These patterns align with known mechanotransductive principles: flatter and deformed cell bodies transmit mechanical force on the nucleus, stretching its pores and enabling the flow of YAP into the nucleus.^[49,50]^ Crucially, Micellangelo preserved these trends, reflecting its ability to emulate biological mechanisms.

**Figure 7.**
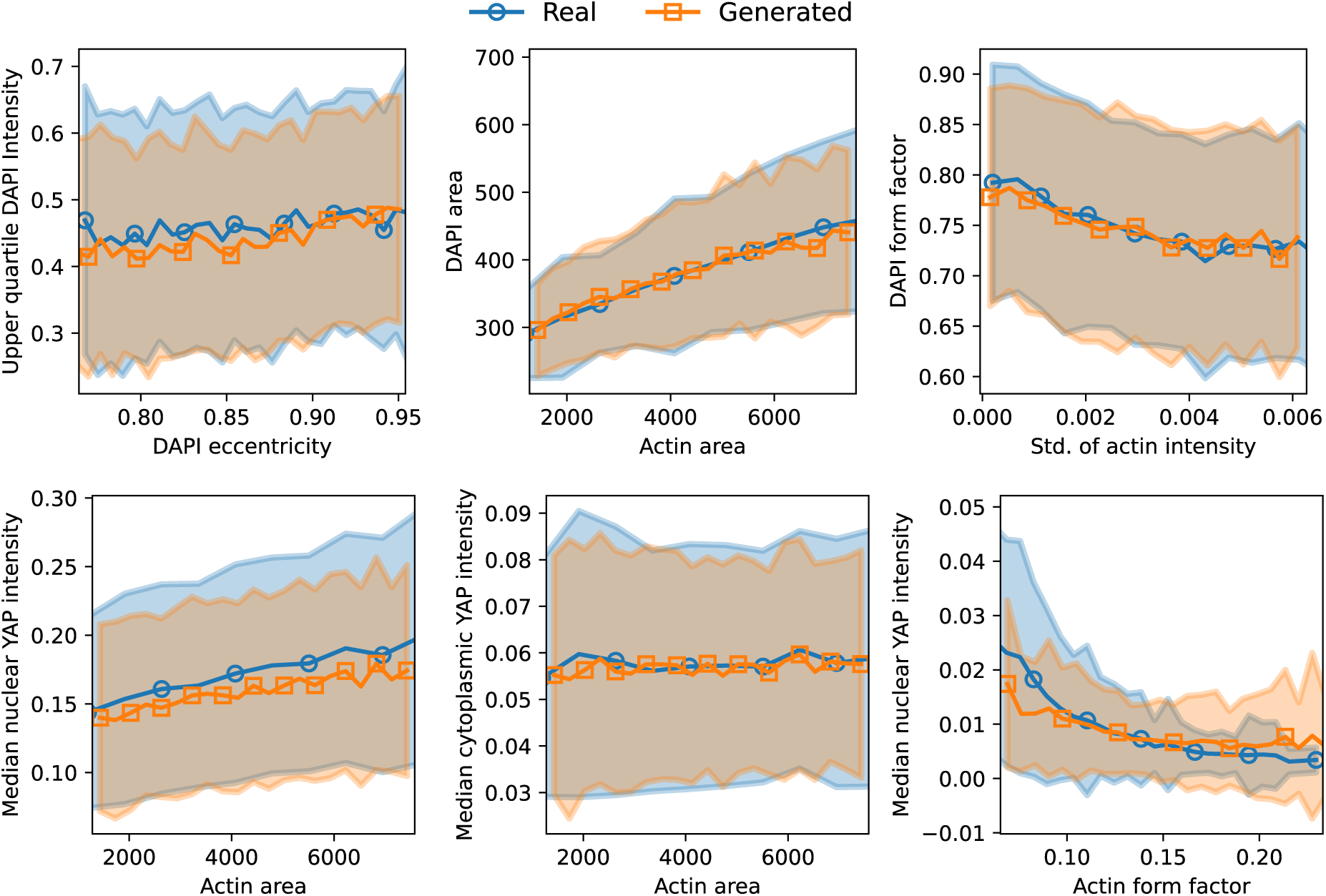
Relationships between biological metrics in real and generated cells. Metrics are calculated on a cell-wise basis; lines and markers indicate means, and the shaded area indicates a single standard deviation.

### 2.5. Structural Consistency Across Topographies

Having analyzed biological metrics at the level of individual cells, we studied the effect of the topography on cells in more detail. To do so, we computed the means of key morphological metrics of the cell body, which are indicative of cell function,^[16]^ across all cells on each topographical insert. This allowed us to assess the averaged morphological response to each surface, regardless of the specific positioning of a cell in the confinement. We then compared these average values between real and generated data, calculating the correlation across surfaces for each metric. Core morphological metrics exhibited high correlations between real and generated data, suggesting that Micellangelo captures the broader trends in how topography shapes cell morphology **(Figure 8, Supplementary Figure S6)**. Features related to cell shape, e.g. extent, eccentricity, orientation, and solidity exhibited the highest correlations, meaning that the effect of the topographical structure on cell shape is captured very robustly. Area and perimeter exhibited slightly lower, but nevertheless strong correlations, suggesting that there is still room for improvement in modeling cell sizes in relation to topography. Overall, most quantities extracted by CellProfiler exhibited strong correlations between the effects of surfaces on real and generated cells.

**Figure 8.**
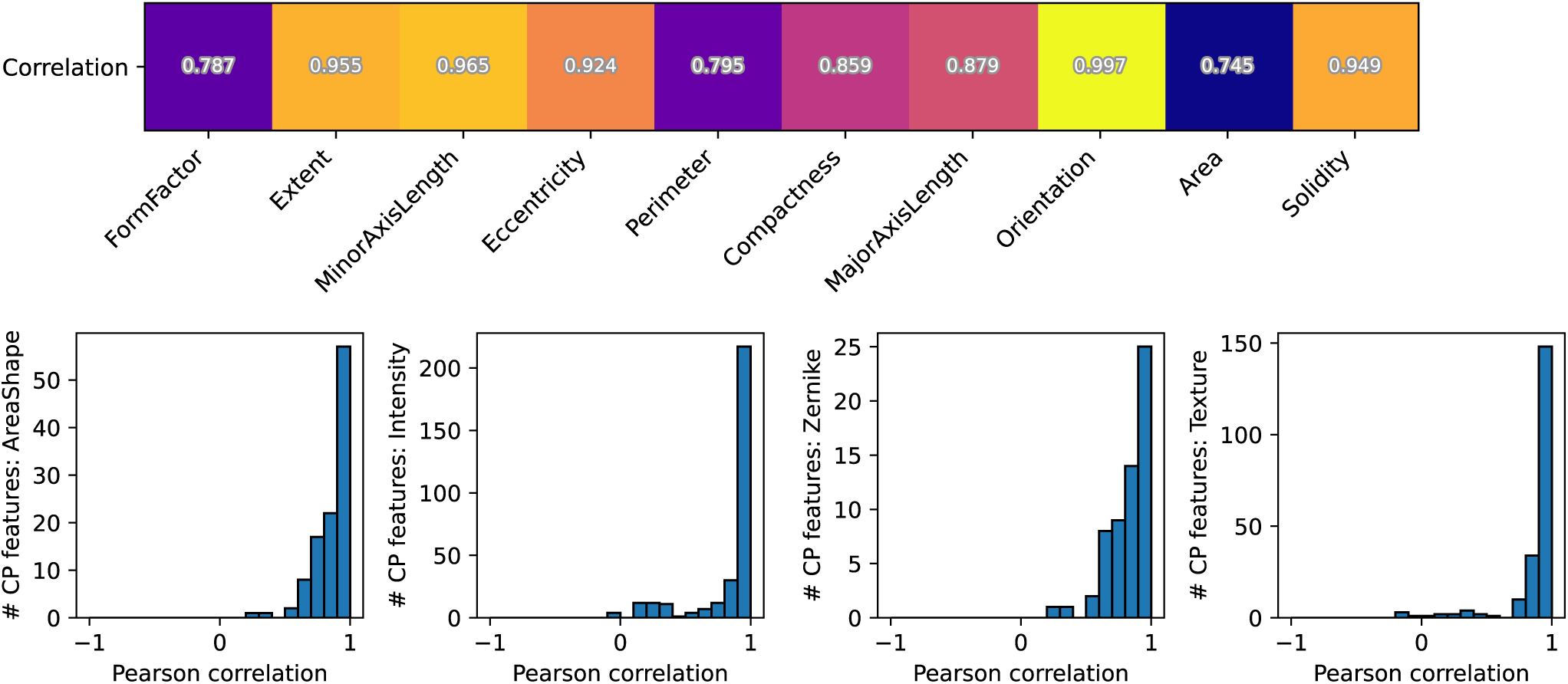
Correlation between real and generated cell morphological features. Top: Pearson correlation between real and generated averaged (insert-wise) morphological metrics; a high value implies that Micellangelo captured the topography-induced morphological effects. Bottom: Histograms of Pearson correlations, calculated as in the top panel, but for all metrics within four categories of CellProfiler features: 110 AreaShape features, 332 Intensity, 60 Zernike, and 208 Texture.

### 2.6. Dose response relationship of biomaterial properties on cell phenotype

To explore potential applications of Micellangelo for in silico cell experimentation and gaining new insights, we conducted two proof-of-concept experiments. These experiments should be viewed as a tool to generate hypotheses and gain inspiration; validation of these hypotheses through wet lab experiments or by other means remains crucial.

First, we studied the effect of the parameter *w* defining the strength of the classifier-free guidance of Micellangelo (see Materials and Methods). Intuitively, a value of *w=0* means that the topographical background is completely ignored in the cell generation process, and the generated cell is basically such that it could have been found on any topography in the training data, while *w=1* indicates that the topography is taken into account in the cell generation process as per usual. Using a biochemical analogy, varying *w* and measuring how generated cells respond could loosely be viewed as characterizing the *dose-response* relationship for topographies. For example, one could think of a gradient of pillar height or material stiffness as physical parameters relating to *w*. Although a useful analogy, we emphasize that *w* does not have a direct interpretation as such a physical mechanism, but we envision that it can still serve as a parameter to investigate how generated cells vary as the strength of the topography-induced conditioning is increased.

When varying *w*, we observed that cells generated under *w=0* do not relate to the topographical landscape, which is evident from e.g. the overlap of the cells with topographical features **(Figure 9A)**. As we increased the value to *w=1*, we noticed that the cells slowly became more realistic for the specific topographical background and morphologically aligned with the landscape surrounding the cell (Figure 9A). Similar to Figure 5, we also measured distributions over morphological features for varying *w* and compared them to the ground-truth distributions. For *w*≤0.5, the distributions showed poor alignment, while w=0.9 and w=1 showed strong alignment as expected **(Figure 9B)**. Finally, we plotted histograms of Pearson correlations between real and generated morphological fingerprints, as explained in Figure 8. For *w=0*, we observed correlations distributed symmetrically around 0.0, indicating that there is no correlation between real and generated morphological features on surfaces; as we increased *w* to 1, the distributions become strongly skewed, and almost all correlations were strongly positive, indicating generated cells that are morphologically similar to real cells **(Figure 9C)**.

**Figure 9.**
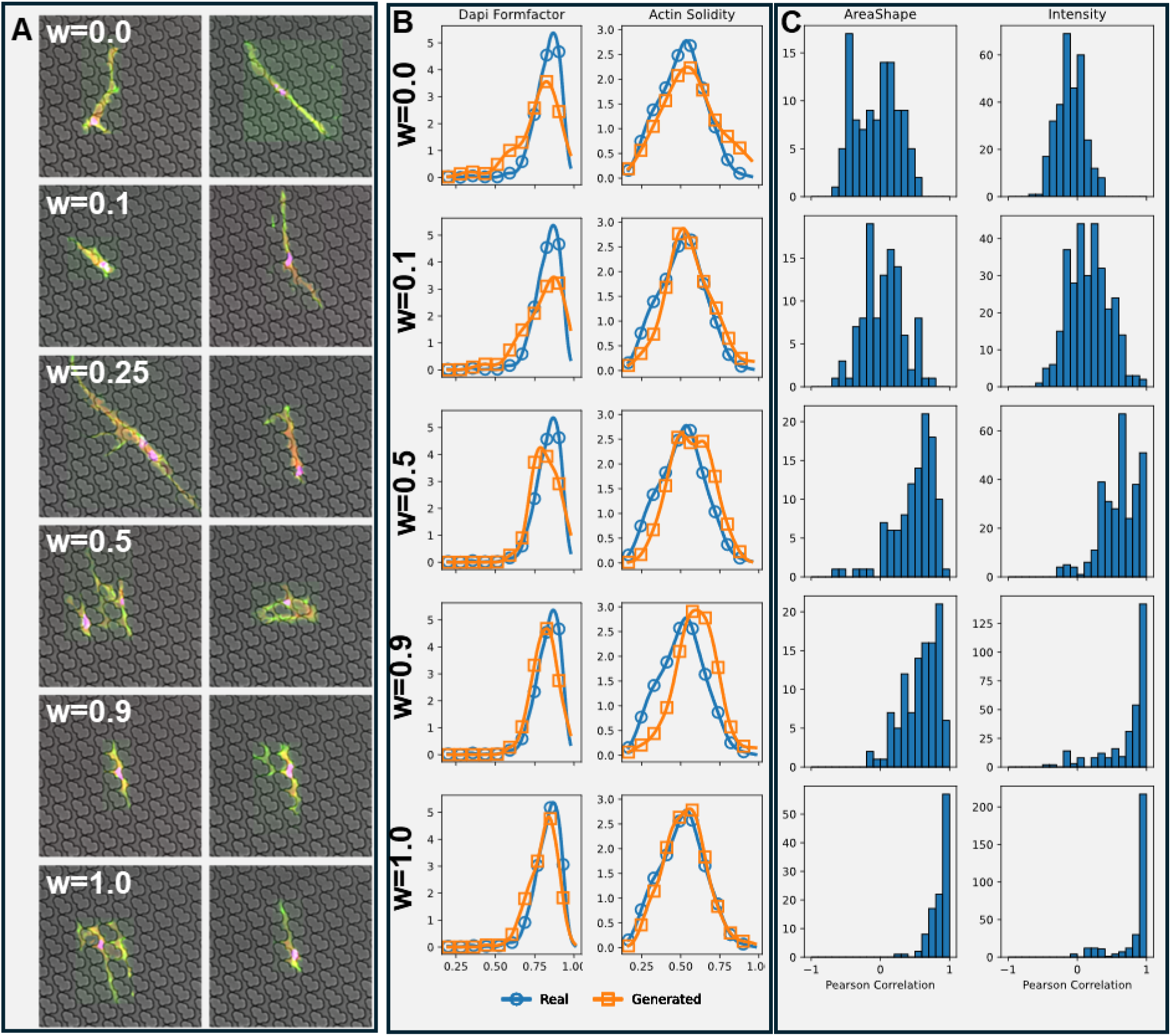
In silico dose–response and validation of topography-conditioned cell generation. (A) Representative generated fibroblast morphologies across conditioning strengths (*w* = 0.0 – 1.0). Lower *w* values correspond to weak conditioning, producing cells that ignore the underlying topography, whereas higher *w* values result in stronger alignment and elongation along topographical features. (B) Distributional comparison of key morphological descriptors between real (blue) and generated (orange) cells on surface 1130. The alignment of DAPI form factor and actin solidity distributions improves with increasing *w*, reflecting the growing influence of surface topography on generated morphologies. (C) Pearson correlations between real and generated morphological fingerprints (110 AreaShape and 332 Intensity features) across surfaces for each *w* value. Correlation strength increases with *w*, indicating enhanced topography-dependent realism of simulated cells.

### 2.7. In silico single cell perturbation experiments

We explored how Micellangelo could be used to perturb cells. We considered a scenario where a cell of a specific shape is defined to be of particular interest. As there is a large variety in cell morphology, this specific shape will be observed only once, even in the case of large-scale screening. Still, even if we observed only one realization of DAPI and YAP stains for any specific phalloidin stain, multiple other realizations could have been plausible. By giving the phalloidin stain as additional input to the model, we were able to investigate the variety of possible corresponding remaining stains for that specific cell shape, similar to,^[51]^ which enabled us to investigate, for example, how YAP responds to nuclei moving around inside the cell body. Such an investigation is relevant to cases where cell function is substantially affected by their shape, which requires specialized experimental setups to investigate.^[52]^ Instead, an AI-driven approach would allow for conducting such experiments in silico, relying on data of cells of various shapes. To facilitate the experiment, we modified the flow matching formulation of our model to have a prior based on Data-Dependent Couplings (see Materials and Methods) and retrained the model with this prior.^[53]^

We considered both phalloidin and DAPI **(Figure 10)** as observed stains. Given an observed phalloidin stain, the nucleus can be found at various locations inside the cell, and YAP intensities varied and tended to be highest around the nucleus (Figure 10). This suggests that not only the observed DAPI and YAP stains were realistic for this particular cell, but the cell could have also been in various other internal states. When considering the DAPI stain as input, we noticed an even greater variety, as the nucleus does not constrain the cell shape and organization as strongly as the cytoskeleton. Consequently, we observed many different realistic cell shapes and corresponding YAP intensities.

**Figure 10.**
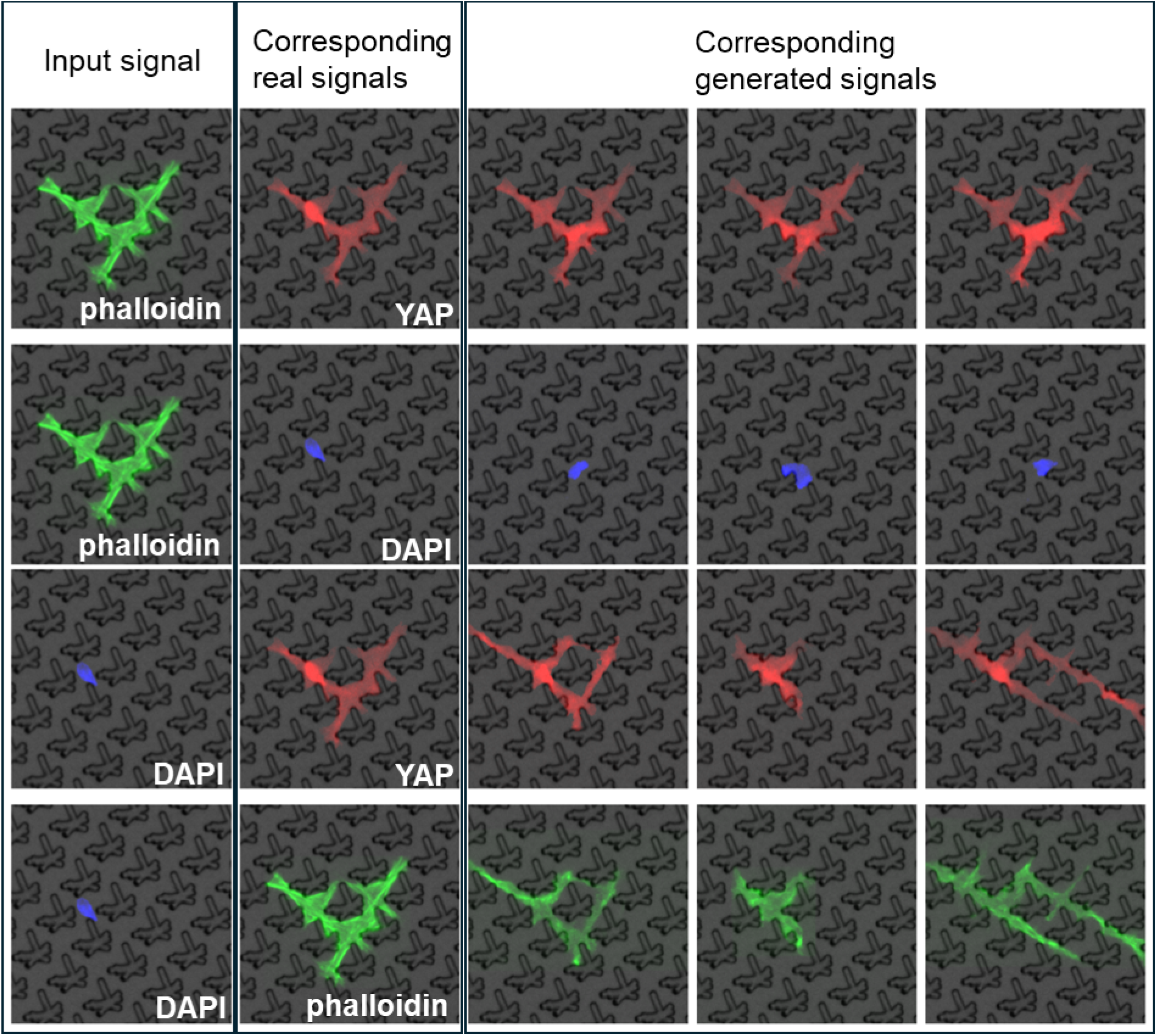
Cross-channel stain generation from partial fluorescence input. Examples of Micellangelo-generated fluorescence channels conditioned on partial input data. Each row corresponds to a specific staining combination, with the leftmost column showing the provided input channel and the remaining columns displaying observed and generated outputs for the missing channels. The model successfully reconstructs nuclei (DAPI, blue), actin filaments (phalloidin, green), and YAP localization (red) consistent with the topographical background. These results demonstrate the model’s ability to infer biologically plausible subcellular organization from incomplete image information.

### 2.8. Graphical user interphase for Micelangelo

To further demonstrate the potential of Micellangelo as a platform for virtual experimentation, we developed a prototype graphical user interface (GUI) that enables a new way of conducting *in silico* experiments within the limitations of the model **(Figure 11)**. This interface provides an interactive environment where users can iteratively design, execute, visualize, and analyze virtual experiments without requiring deep technical expertise in generative modelling or computer programming.

**Figure 11.**
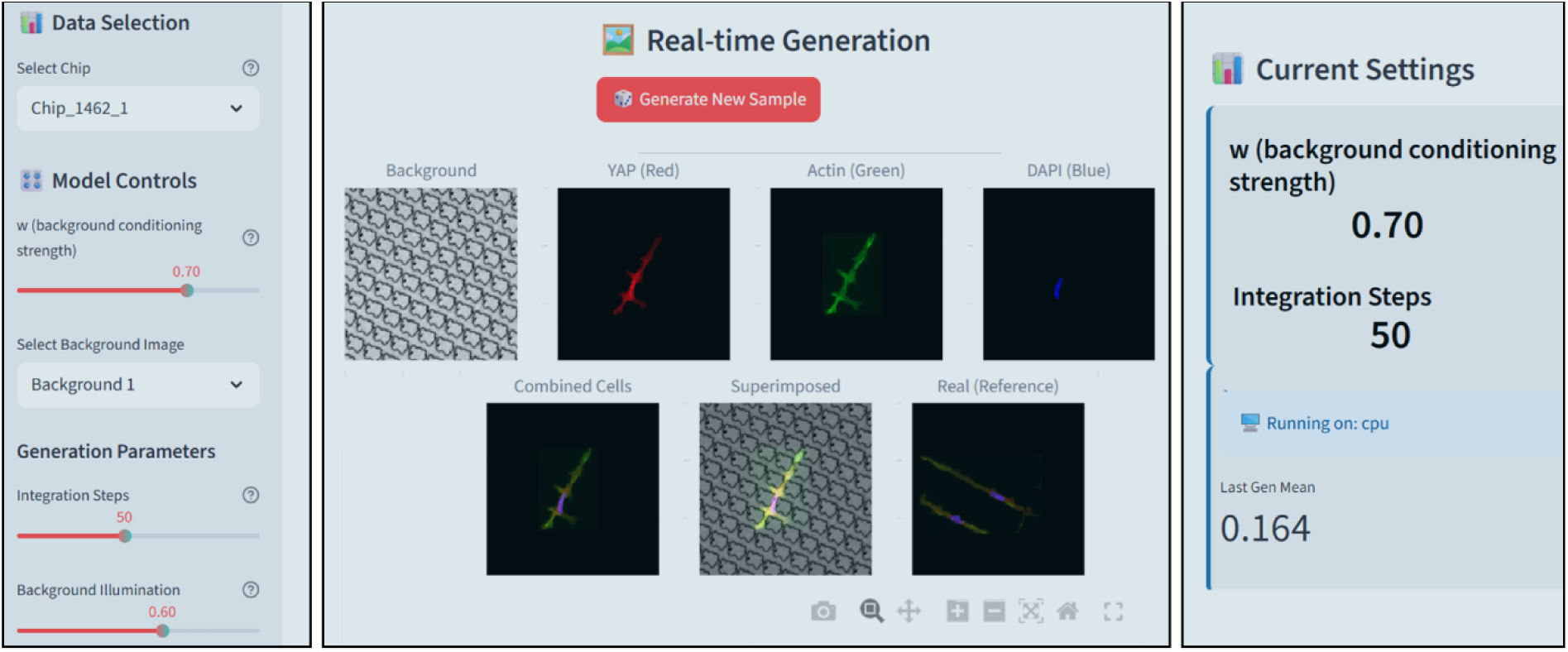
Graphical user interface (GUI) for *in silico* experimentation with Micellangelo. Prototype version of the Micellangelo GUI demonstrating an interactive workflow for designing, running, and analyzing *in silico* cell–material experiments. The interface consists of three main components: (Left) Input and model control panel, where users can select two surface topographies, adjust the background conditioning strength *w*, and fine-tune image generation parameters such as integration steps and illumination; (Center) Real-time visualization panel, which displays generated fluorescence images for individual channels (YAP, actin, DAPI), combined overlays, and superimposed results alongside real reference images, with interactive inspection tools such as zoom and pan; and (Right) Dynamic analytics panel, where current model parameters, system performance, and real-time calculated metrics are displayed. Together, these components enable users to iteratively explore how topographical cues influence cellular morphology within the defined limitations of the model.

The GUI is structured into three main functional blocks. The first block allows the user to configure key parameters of the *in silico* experiment, including the selection of two distinct surface topographies, adjustment of the background conditioning strength *w* (as described in Figure 9), and control over image generation parameters such as the number of integration steps and background illumination. By modifying these settings, researchers can probe how variations in topographical cues influence cellular morphology and mechanotransductive responses.

The second block provides real-time visualization of generated results. Users can observe synthetic fluorescence images for individual channels (YAP, actin, and DAPI), view combined overlays, and compare them directly to real reference images. The interface also includes standard tools, such as zoom and pan, which facilitate detailed inspection of generated cellular morphologies and enable direct qualitative comparisons between different conditions.

The third block presents real-time quantitative analysis and exportation of the generated data. As images are produced, morphological and intensity-based metrics are calculated and displayed as statistical plots. This immediate feedback allows users to rapidly assess trends, compare outcomes between conditions, and iteratively refine experimental parameters. In addition to on-screen visualization, the GUI also provides an option to save and export the generated data, including both raw images and computed metrics, enabling further downstream analysis or integration with external data processing pipelines.

Together, these components establish an interactive workflow that integrates design, visualization, analysis, and data export within a single platform. Although the current version of the GUI is limited to the surfaces and features available in the training data, it provides an important first step towards integrating generative models into biomaterials research workflows and lowering the barrier for using AI-based *in silico* experimentation. The prototype version of the GUI is publicly available on GitHub. ^[54]^

## 3. Discussion

In this study, we presented Micellangelo, a generative model capable of simulating realistic, high-resolution fluorescence staining images of single cells cultured on topographies. To train Micellangelo, we introduced a new dataset of high-resolution images of fibroblasts stained for DAPI, phalloidin, and YAP. Qualitative results indicated that simulated cells are visually convincing and similar to real cells, whereas quantitative results based on image analysis demonstrated alignment with real cells in terms of topography-induced and biologically relevant metrics such as area, eccentricity, and YAP localization. Going beyond cell image simulation, we took the first steps towards in-silico experimentation: first, we varied the strength of the topographical conditioning and analyzed its effect on cell morphology. Second, we explored a proof-of-concept method to investigate how cells respond to perturbations, for example, how YAP intensities change as the nucleus is positioned in different parts of the cell body.

Our results align with recent works on generative modeling for cell imaging experiments. (Bourou, Mahanta, et al., 2025b; Bourou, Segade, et al., 2025b; Bourou Anisand Boyer, 2024b; Donovan-Maiye et al., 2022c; Lamiable et al., 2023b; Zhang et al., 2025b) These studies have shown that generative models are capable of generating realistic cell images, even when conditioned on scalar variables representing, for example, chemical perturbations. Compared to these works, we adapted the modeling machinery for the topographical biomaterials setting, which requires conditioning on image-like inputs rather than scalar values or encodings of the material composition. Moreover, we explored how such a model could be used to potentially conduct single cell perturbation experiments in silico. Overall, this work aims to introduce technological state-of-the-art and its capabilities to the biomaterials community and to inspire new research directions on in silico experimentation. It is important to note that generative models in general express relationships that are correlative, rather than causal, and observed phenomena may even be the result of hallucinations. As such, it remains imperative to verify any in silico observations in laboratory experiments before the results can be trusted and integrated in the scientific body of knowledge. In this light, we envision Micellangelo and other generative models as a tool to assist cell biologists in formulating research questions, generating hypotheses, and designing laboratory experiments. We consider our work a proof-of-concept that generative AI can serve as a digital twin of laboratory experiments and aid in the investigation of cellular responses to biomaterials. To verify this approach, we focused on an experiment with manageable biological variability: we considered ten topographies of polystyrene substrates, single fibroblasts, one passage number, and DAPI, phalloidin, and YAP staining. Our data acquisition protocols enable future studies that consider a larger variety of material properties, such as chemical composition and more diverse topography designs. Moreover, the modeling approach straightforwardly generalizes to additional fluorescent stains, and can be extended to support different cell types and material compositions by encoding this information and providing it as an additional conditioning signal to the model.

Currently, developing and using generative models requires specialized expertise. Low barrier-to-entry software, akin to CellProfiler,^[61]^ could democratize the technology and empower cell biologists to explore the possibilities of in silico experimentation. Standardized and open data-sharing practices within the biomaterials community are crucial to facilitate the development of such software and ultimately progress towards more general and widely applicable in silico biomaterial experiment models.

## 4. Materials and Methods

Detailed experimental procedures, including topography fabrication, cell culture, fixation and staining, fluorescence imaging, segmentation, and quality control, are provided in the **Supplementary Materials and Methods**. Below, we summarize the computational framework underlying the Micellangelo model.

### 4.1. Generative model

We designed Micellangelo using the *flow matching* framework,^[32]^ and we based our implementation primarily on ^[43]^. We briefly summarize this approach here, and refer to for technical details.^[43]^ The core of the approach entailed designing a neural network that models a continuous transformation (called a *normalizing flow*) from a prior distribution *p*_0_ to the distribution *p*_1_ over cell images. As is common practice, we chose *p*_0_ to be a standard Gaussian. To train the model, rather than using the original max-likelihood objective of,^[32]^ the neural network *v*_*θ*_(*x*_*t*_, *t*) was trained against a conditional time-dependent vector field *u*(*x*_*t*_, *t*∣*Z*) as specified in^[44]^, where *Z* ∼ *q*(*Z*)^[32]^; here, *t* denotes an internal fictive time variable used by the model to condition the vector field on. We chose *Z* = (*x*_0_,*x*_1_) to be a pair of samples from prior and data distribution, and *q*(*Z*) = *π*(*x*_0_,*x*_1_) as the optimal transport map *π* between *p*_0_ and *p*_1_, following Tong et al.^[43]^ To condition on the topographical background *C*, we employed classifier-free guidance, where *VṼ*_*θ*_(*x*, *t*∣*C*) = (1 ― *w*) ⋅ *v*_*θ*_(*x*_*t*_, *t*) +*w* ⋅ *v*_*θ*_(*x*_*t*_, *t*∣*C*). After training, synthetic data was simulated by numerically integrating the vector field 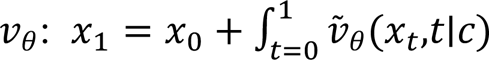, where *x*_0_ ∼ *p*_0_, following Chen et al.^[62]^

### 4.2. Neural network architecture

Following the original work and its recent improvements of ^[63,64]^ we implemented a UNet neural network architecture to parameterize *v*_*θ*_. As opposed to scalar conditioning, we relied on the topography of the material as a conditioning signal for generating cell images, which is in itself an image. As a whole, the model processes this input image of the topography together with the noisy image of the cell *x*_*t*_ to produce a vector that points in the direction to ‘denoise’ the image towards a realistic cell image free of noise. Specifically, our implementation used a separate topography embedding network *Φ*, which we trained in end-to-end fashion together with *v*_*θ*_*C*^′^ = *Φ*(*C*) is then passed as input to *VṼ*_*θ*_(*x*_*t*_, *t*∣ *C*′). The design pattern of *Φ* aligned with the UNet, as it consists of convolutional blocks followed by max pooling layers. The resulting embeddings have the same spatial scales as in the UNet architecture. We integrated the conditioning on *C*′ in the UNet through additive composition of the embeddings and the internal UNet embeddings at each spatial scale, similar to how conditioning on scalar features is usually done in generative computer vision models. The UNet starts with 32 channels and consists of 5 down sampling layers, where the number of channels doubles only in the last three down sampling layers, and the corresponding up sampling layers. Self-attention with 4 heads is applied only in the bottleneck layer. We trained the model for 24 hours (145000 training steps) using the Adam optimizer with a learning rate of 0.0002, a dropout rate of 0.1, EMA with a decay of 0.9999, gradient norm clipping with a maximum of 1.0, and a batch size of 32. We also segmented and masked the F-actin and DNA channels to remove background illuminance and cropped images to the center 300×300 pixels when providing training samples to the model. Further architectural and training implementation details can be found in our code https://github.com/cbite/Micellangelo.

### 4.3. Cell stain completion with data-dependent couplings

To facilitate cell image generation with partially observed stains, we retrained the model using a prior distribution defined through data-dependent couplings.^[53]^ Specifically, we followed the exact procedure as described above, with the difference that zero, one, or two out of the total three cell imaging channels are observed at random with probabilities 0.1, 0.1, and 0.8 respectively; the unobserved channels were Gaussian noise as above. Further, the model was provided with a boolean mask indicating which channels are observed as additional input. During inference, the model received a single stain of a cell as input following the same procedure as during training, so that it generates a plausible completion of the remaining two channels.

## Acknowledgements

The authors would like to acknowledge Sander Derwig, whose master’s thesis work served as an inspiration and foundation for the analysis workflow developed in this study. We are grateful to the Cell Culture Facility at Eindhoven University of Technology, under the management of Marloes Janssen, where all cell experiments were performed. We also thank the Department of Biomedical Engineering at Eindhoven University of Technology for providing the opportunity to present our work during the REM Cluster Colloquiums, which helped shape the direction and finalization of this manuscript. Lastly, we acknowledge the creators and community of CellProfiler for enabling robust and reproducible image analysis. During the preparation of this work, the authors used **ChatGPT (OpenAI)** for language refinement, manuscript structure optimization, and guidance on compliance with journal submission guidelines. After using this tool, the authors carefully reviewed and edited the content as needed and take full responsibility for the accuracy, completeness, and integrity of the published article.

## Data Availability Statement

The raw microscopy data that support the findings of this study is available from the corresponding author upon reasonable request.

Preprocessing pipeline and the model are available online https://github.com/cbite/Micellangelo/

The current alpha version of the GUI is publicly available on GitHub at https://github.com/cbite/Micellangelo/tree/main/GUI/

Received: ((will be filled in by the editorial staff))

Revised: ((will be filled in by the editorial staff))

Published online: ((will be filled in by the editorial staff))

## Supporting Information

## 1. Results

### 1.1. Database of single cell images on topographies

**Supplementary Figure S1.**
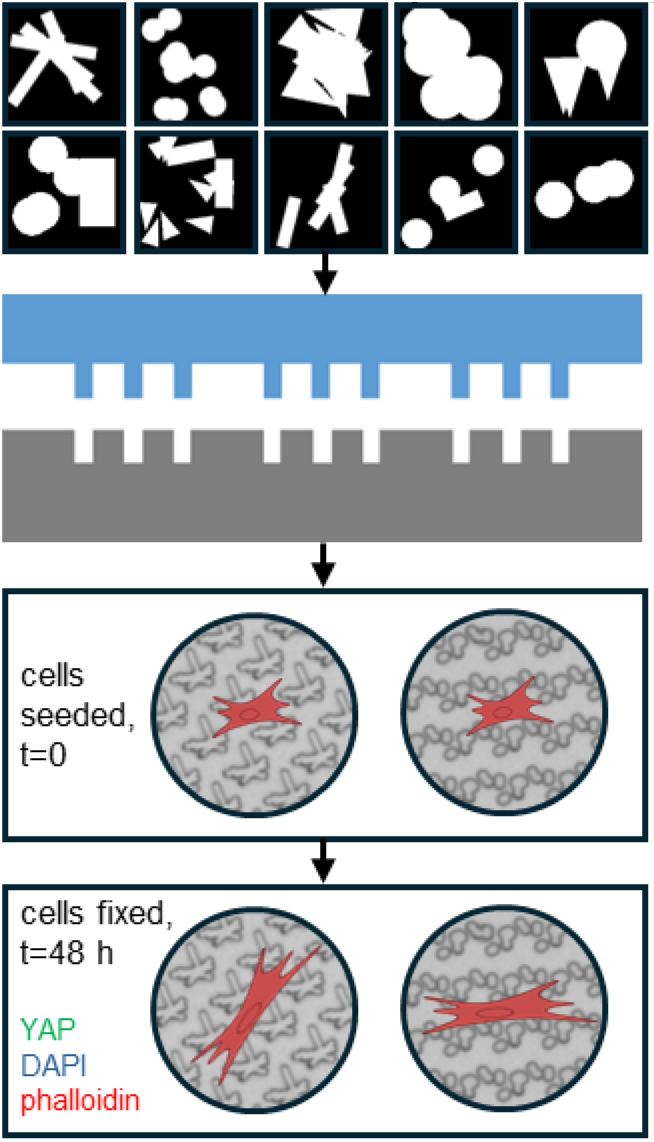
Experimental outline to generate the image library. Experimental workflow to generate a high-quality dataset of fibroblasts cultured on engineered micro-topographies. A selection of 10 previously validated surface designs (top row) was fabricated into 13 mm diameter polystyrene (gray) inserts using silicon molds (blue). Fibroblasts were cultured for 48 hours and then stained for three markers: DNA (DAPI, blue), the mechanosensitive transcriptional regulator YAP (green), and the actin cytoskeleton (phalloidin, red). Fluorescence imaging captured how surface micro-topography modulates cell morphology and subcellular organization.

**Supplementary Figure S2.**
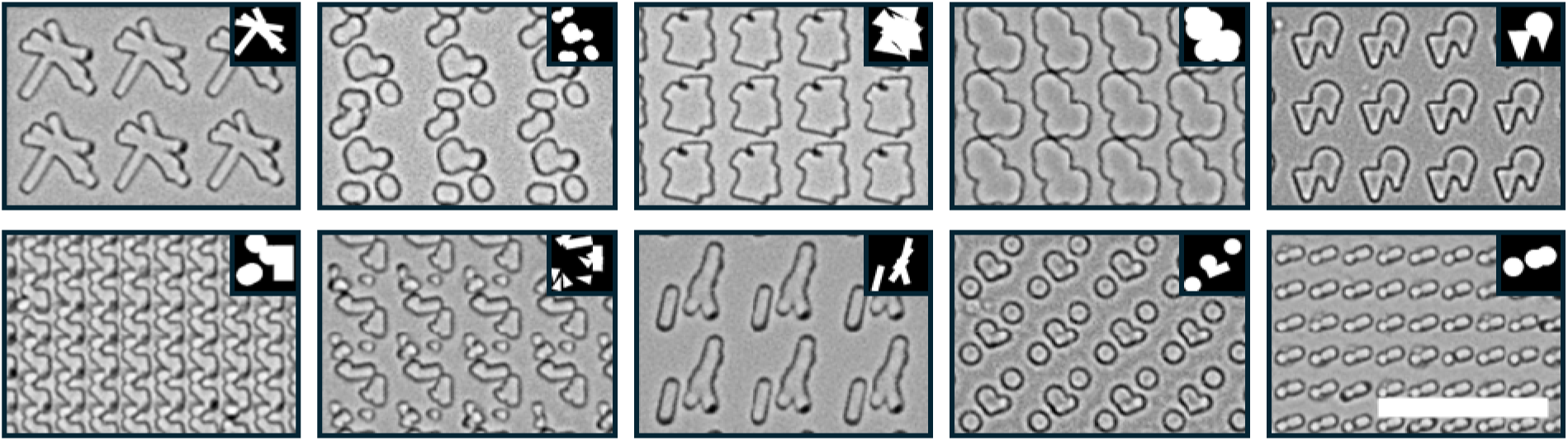
Brightfield images of surface inserts used as substrates for cell culture experiments. Each panel shows a surface topography insert with its corresponding feature idx: top row (left to right): 1130, 1330, 1462, 1476, 1689; bottom row (left to right): 1710, 1913, 2065, 2088, 2113. Black and white inserts display the pattern generated by the design algorithm. Scale bar: 50 µm.

### 1.2. Segmentation and object extraction for single-cell quantification

To enable quantitative analysis of fibroblast morphology across topographically patterned surfaces, we developed and applied an intensity-based segmentation pipeline using *CellProfiler*. The goal was to identify individual cells and their subcellular components as discrete objects, extract relevant features, and preserve spatial context relative to the substrate. For each of the ten topographies, a set of 50 representative images was selected and processed to calibrate segmentation parameters. Illumination correction was first applied using a background subtraction method for the DAPI channel and a regular correction function for YAP and phalloidin. To suppress imaging noise and improve object boundary detection, a Gaussian smoothing filter was used with kernel sizes matched to expected object diameters: 80 pixels (∼45 μm) for nuclei and 250 pixels (∼140 μm) for cytoplasmic markers.

Next, segmentation thresholds were set manually per topography and per channel (DAPI, phalloidin, YAP), informed by suggested ranges from the *Measure Image Quality* module. Nuclei stained with DAPI were defined as primary objects and constrained to an accepted size range of 15–40 pixels in diameter (∼8–22 μm). Cytoplasmic objects stained with phalloidin and YAP were then segmented as secondary objects associated with each nucleus. To ensure data quality, all touching or overlapping objects, as well as those intersecting image borders, were systematically excluded. Thus, only isolated, non-touching single cells were retained for downstream analysis.

For each segmented object, we extracted 73 intensity and morphology features, for later analysis. The segmentation outlines were mapped onto the corresponding brightfield image to define spatial alignment with the underlying topography **(Supplementary Figure S3)**.

**Supplementary Figure S3.**
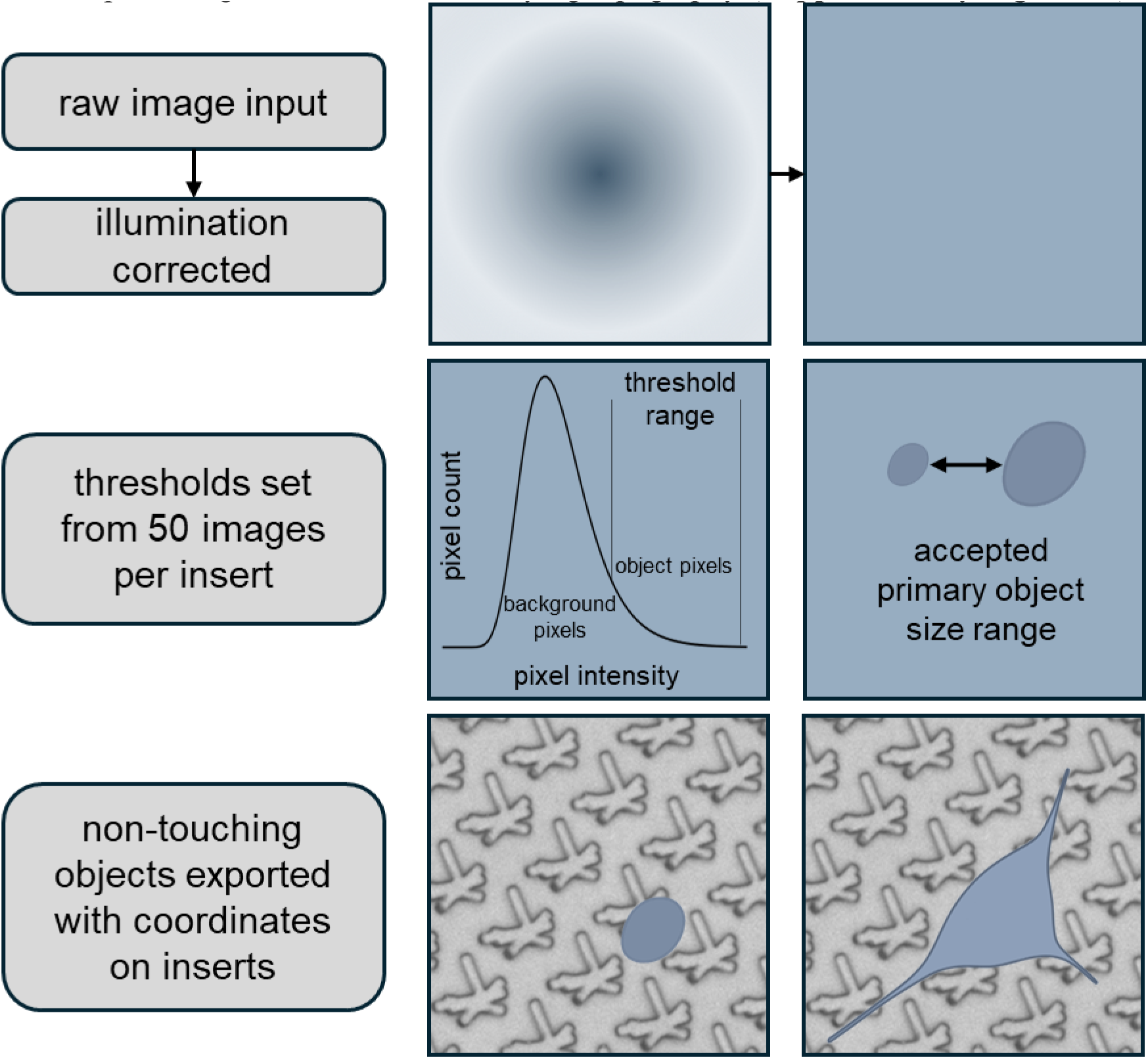
Workflow to segment cells and define their spatial location. Images underwent illumination correction. For each insert, 50 representative images were visually inspected to define thresholds for primary object intensity and size. Only non-touching objects within the accepted size range and above the intensity threshold were segmented. The outlines of segmented signals were then exported along with their spatial positions on the brightfield insert images. The graph indicates how intensity thresholds were defined, while the dot size schematic shows the accepted object size range.

Across all surfaces and technical replicas, a total of 17,852 single nuclei were segmented prior to quality filtering. However, this total was disproportionately influenced by surface 2065, which alone accounted for 6,755 segmented objects. Upon inspection, this surface displayed a recurring DAPI staining artifact of numerous small, bright specks lacking corresponding phalloidin and YAP signals. These specks fell within the acceptable size range and were therefore initially segmented as nuclei, but were systematically excluded during downstream quality control.

When excluding surface 2065 from the analysis, the dataset comprised 11,097 nuclei across the remaining surfaces, yielding an average of 1,010 ± 163 nuclei per surface. This distribution reflects a more accurate representation of object detection across topographies with high-quality staining. The segmentation results formed the basis for subsequent object classification and morphological profiling, described in the next section.

### 1.3. Segmentation quality control and outlier removal

To ensure that we included only correctly segmented cells, we implemented a two-step quality control (QC) process combining classifier-based filtering and morphological outlier detection. First, we trained a dedicated object classifier for each surface using CellProfiler Analyst (CPA). Per surface, 50 well-segmented cells and 50 artifacts were manually labeled to define a training set. The classifier was then used to identify low-quality or mis-segmented objects across the entire dataset, with examples including DAPI-positive specks **(Supplementary Figure S4A)** and debris-laden aggregates not associated with true cells.

Following classifier-based filtering, we applied a second filtering step targeting biologically implausible morphologies based on the nuclear-to-cytoplasmic area ratio (NtoC). We removed cells of which the logarithm of NtoC was beyond 1.5 times the interquartile range from the first and third quartiles. This helped remove segmentation artifacts such as overlapping nuclei, ghost cells, or debris captured by DAPI but lacking a surrounding cell body (Figure 6B).

**Supplementary Figure S4.**
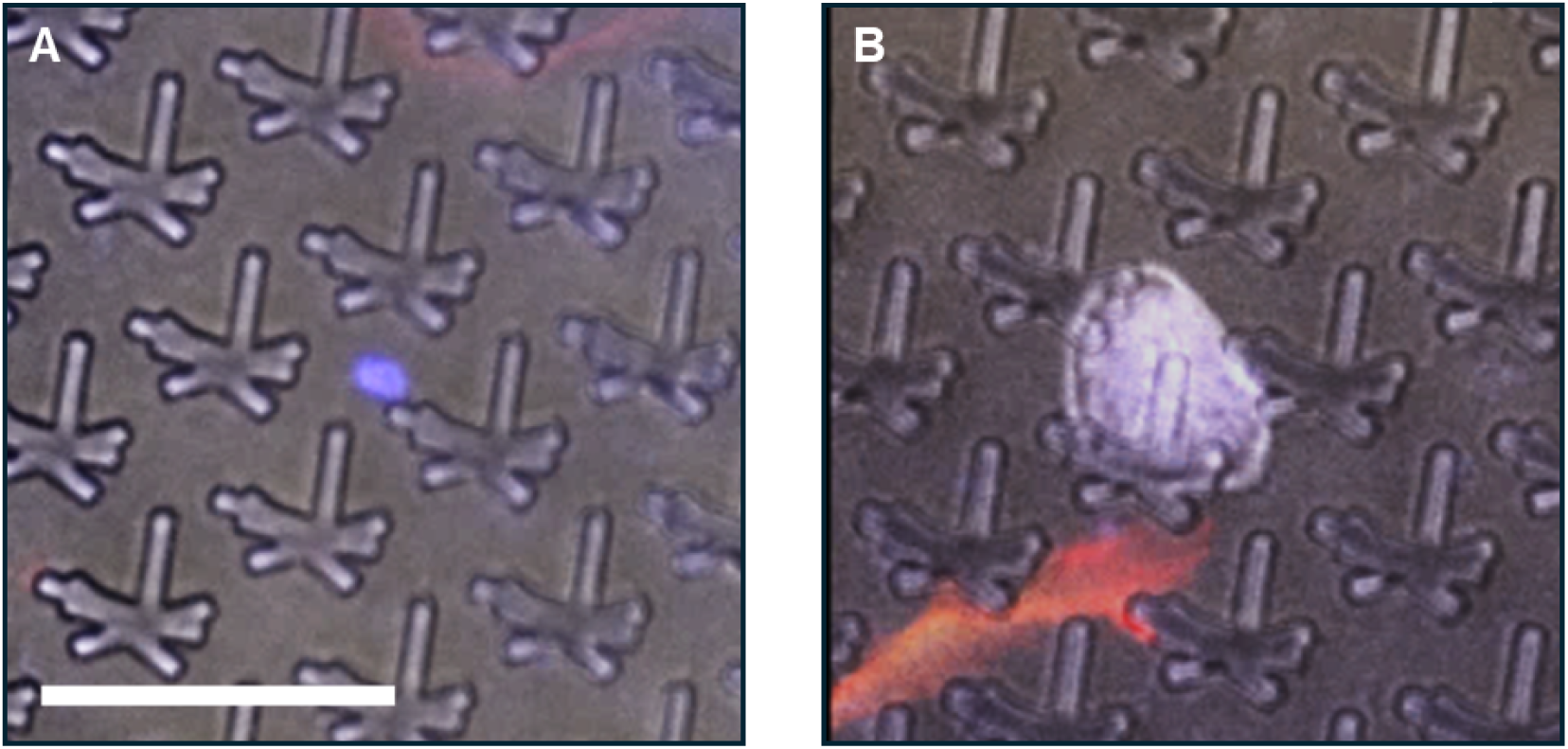
Examples of objects detected by quality control. (A) A DAPI staining artefact was identified during classification in CellProfiler Analyst’s Classifier module, showing a small, bright blue spot not associated with a true nucleus. (B) A piece of debris was flagged as an outlier based on an abnormally high nucleus-to-phalloidin area ratio. Composite image channels: Brightfield (gray), DAPI (blue), Phalloidin (red), YAP (yellow). Scale bar: 50 µm.

This combined QC approach removed low-quality segmentations, particularly from surfaces prone to staining artifacts. For example, surface 2065 alone accounted for 6,147 removed objects due to recurring DAPI speckles, while other surfaces showed markedly lower exclusion rates. The final number of high-confidence cells retained after QC **(Supplementary Table 2)** represents the dataset used for downstream feature extraction and modeling. This robust filtering pipeline ensured that only well-segmented, biologically plausible cells were retained for analysis.

### 1.4. In silico generation of cell images. Realistic single cell generation

**Supplementary Figure S5.**
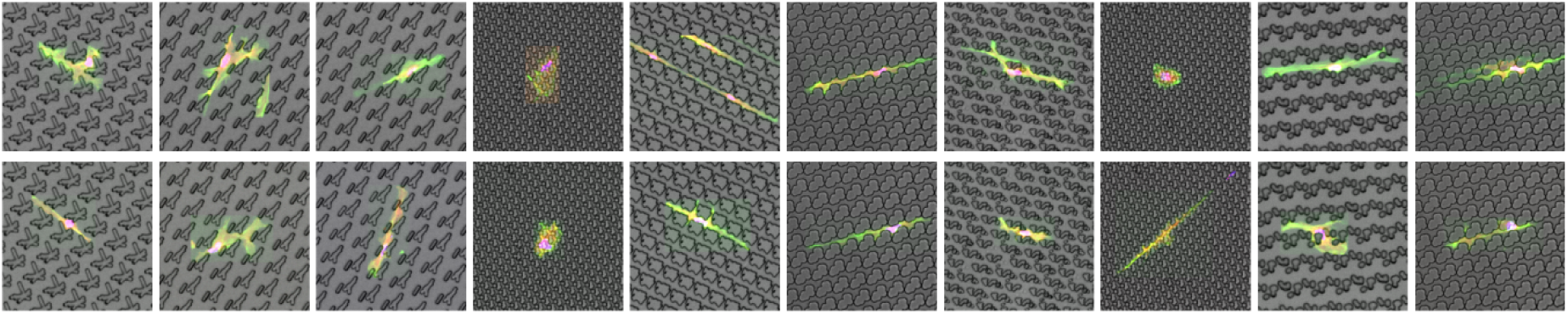
Comparison between real and Micellangelo-generated fibroblast images. The top row displays fluorescence images of real fibroblasts cultured on micro-topographical substrates and stained for nuclei (DAPI), actin (phalloidin), and YAP. The bottom row shows corresponding synthetic cells generated by Micellangelo, exhibiting similar morphological and staining characteristics. Visual alignment between real and generated cells highlights the model’s capacity to replicate biologically relevant cellular features.

### 1.5. Structural Consistency Across Topographies

**Supplementary Figure S6.**
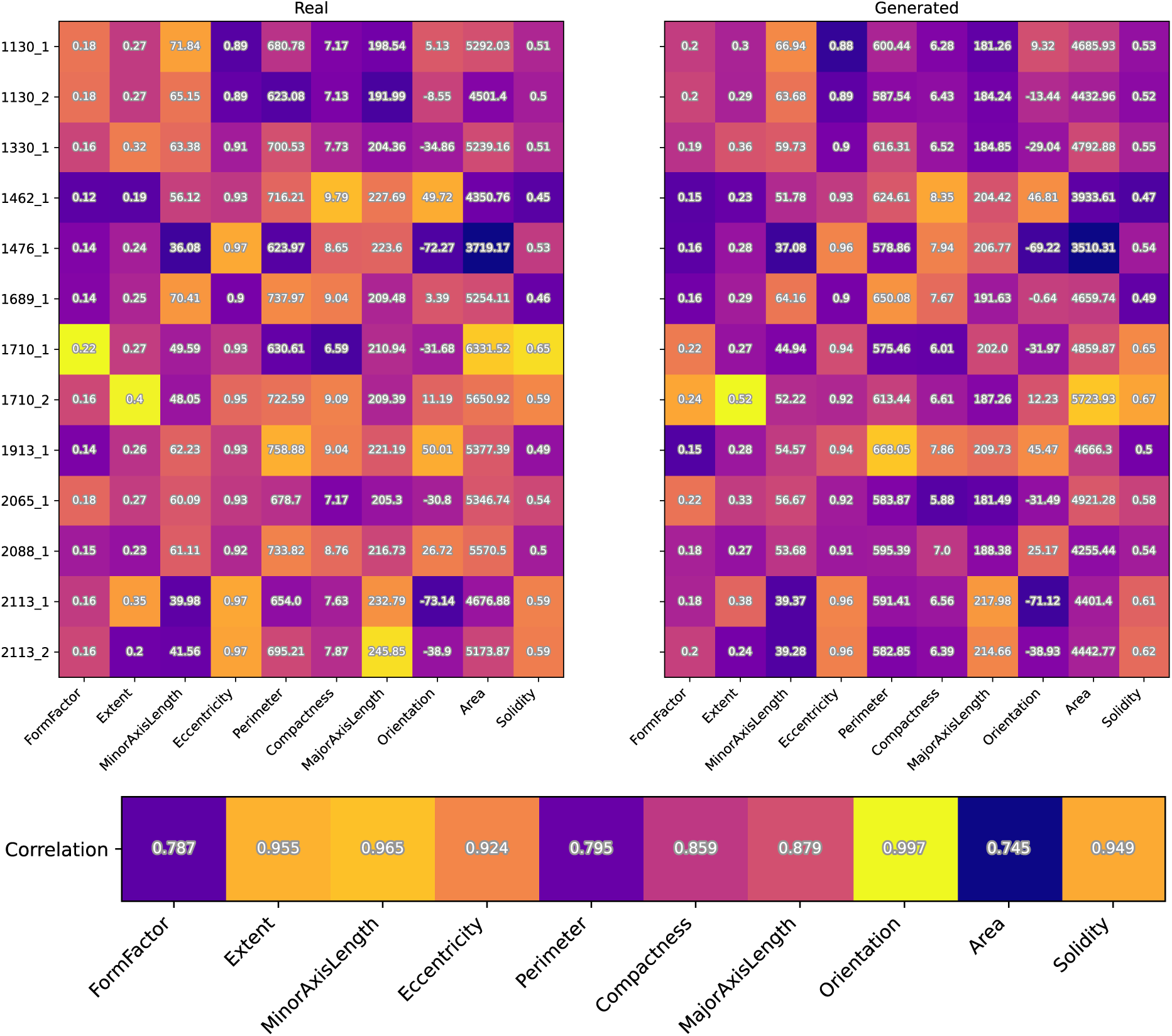
Correlation between real and generated cell morphological features. Average morphological metrics for each surface for real (left) and generated (right) data, calculated as surface-wise averages of morphological metrics. Each row can be interpreted as the ‘morphological fingerprint’ induced by the surface.

### 1.6. Topography-Level Effects and Localized Confinement

When considering relative positioning of nuclei to topographical features, we expect that nuclei in an area of high confinement need to ‘squeeze’ to fit in the spatial confinement, leading to a lower form factor. Conversely, nuclei that are in areas with more space between the confinement are not under mechanical tension and consequently remain more circular, reflected in a high form factor value. We observed this effect for both real and generated cells **(Supplementary Figure S7)**. This suggests that Micellangelo not only captures the general effect of the topography on cellular morphology, but also respects the localized confinement of the cell induced by its topographical surroundings, in line with our qualitative observations of Figure 9b.

**Supplementary Figure S7.**
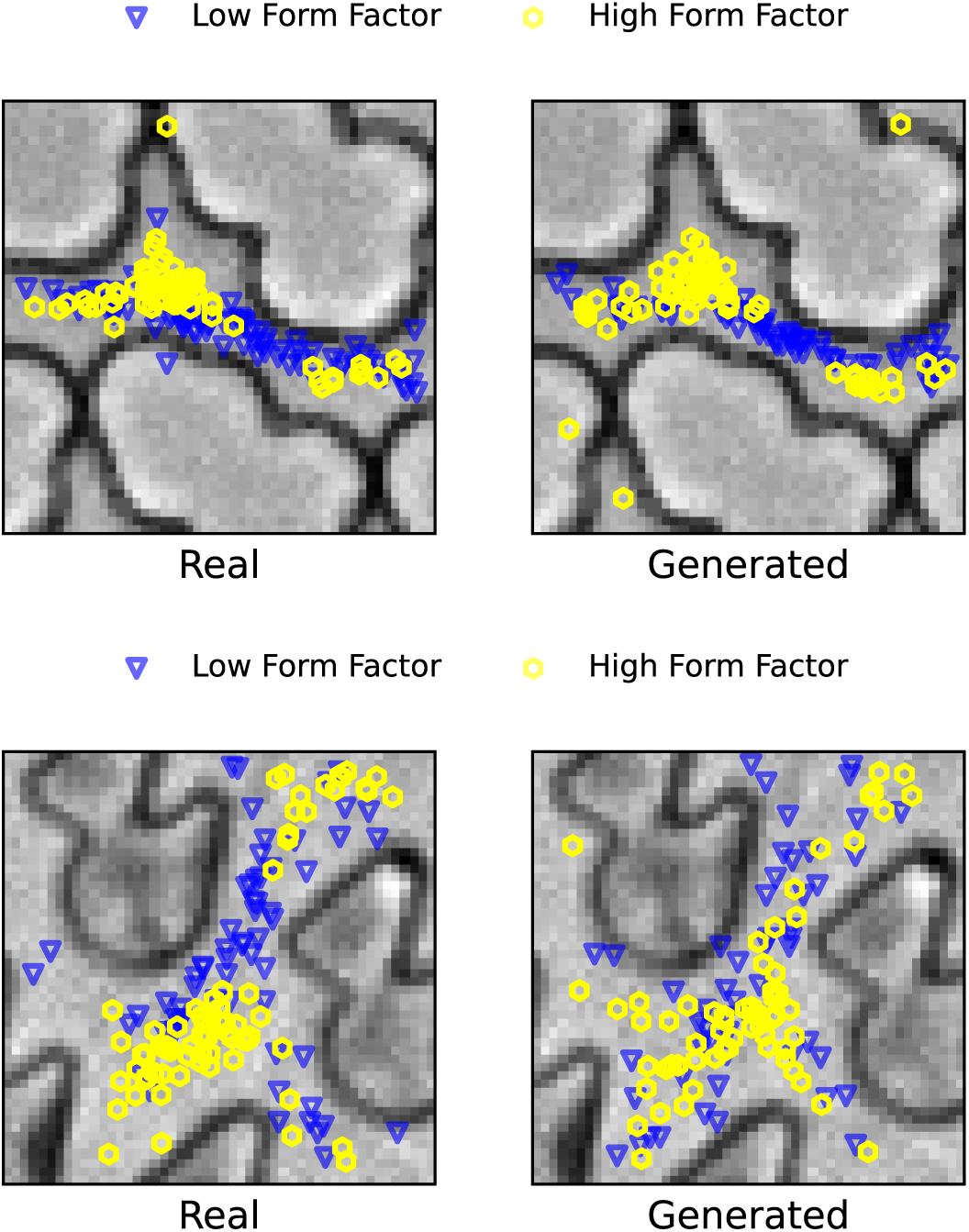
Location-specific nuclear shape in real and generated cells. Cell nuclei centroid locations on surfaces 1476 (top) and 1689 (bottom), for nuclei with form factors in the lowest (blue triangles) and highest (yellow circles) 20% of nuclei form factors. Nuclei with the highest form factors are concentrated in areas with the least stringent confinement on the topographical coordinate space, meaning that nuclei with more space tend to be more circular.

## 2. Materials and Methods

### 2.1. Topography fabrication and quality control

A library of polystyrene substrates with engineered micro-topographies was fabricated using a multi-step hot embossing process adapted from standard TopoChip protocols. In short, negative molds were created by casting polydimethylsiloxane (PDMS; Sylgard 184, Dow Corning) onto etched silicon wafers and cured at 85°C for at least 2 hours. Positive OrmoStamp molds (Micro Resist Technology GmbH) were generated via UV curing (irradiance 15, dose 1100 mJ/cm²) followed by thermal post-curing. Prior to embossing, molds underwent oxygen plasma treatment (0.8 mbar O₂, 100 W, 30 s) using a plasma asher and were subsequently silanized with trichloro(1H,1H,2H,2H-perfluorooctyl)silane in a vacuum desiccator to facilitate demolding and prolong mold usability. Final topographies were embossed into 190 µm-thick polystyrene sheets (Goodfellow) at 140°C under 5T pressure for 10 minutes using a brass–Teflon–Ormomold sandwich in a hydraulic press.

Demolding occurred at 90°C. Surface fidelity was verified using light microscopy at 10× and 40× magnification. Complete fabrication SOPs and material specifications are available at [https://github.com/cbite/TopoChip-analysis/blob/main/1-fabrication.7z].

### 2.2. Surface preparation and sterilization

Embossed polystyrene sheets were manually cut into circular inserts (13 mm diameter) using scissors and positioned into the wells of a sterile 24-well cell culture plate (CELLSTAR, Greiner), topography-side facing up. Inserts were held in place using custom-sized silicone O-rings (Ø = 1.5 cm, ERIKS) to prevent floating during cell culture. To remove residual manufacturing contaminants and ensure reproducibility across batches, O-rings were subjected to 24-hour incubation in sodium dodecyl sulfate (SDS; Sigma-Aldrich; 1% w/v) on a rocking plate at 300 rpm. This was followed by three sequential washes with sterile phosphate-buffered saline (PBS, Sigma-Aldrich). For surface sterilization, ethanol (VWR Chemicals, BDH Chemicals, 500 μL of 70%) was added to each well, ensuring complete submersion of the surface, and surfaces were incubated for 30 minutes in a laminar flow safety cabinet. Ethanol was then aspirated, and surfaces were rinsed three times with sterile PBS (1 mL). After the last wash, PBS was aspirated and plates were used immediately for cell seeding.

### 2.3. Cell culture and seeding

Normal human dermal fibroblasts (NHDF; Lonza, CC-2511) were cultured in Dulbecco’s Modified Eagle Medium (DMEM; Gibco, Fisher Scientific, 42430), supplemented with fetal bovine serum (FBS; Sigma-Aldrich, 10%) and penicillin-streptomycin (Gibco, 1%). Cells were thawed at passage 1 and maintained in T75 flasks at 37°C in a humidified 5% CO₂ incubator for one week prior to the experiment. During expansion, cells were sub-cultured every 2–3 days at ∼80% confluency, with an observed doubling time of 36 to 48 hours. Cells of passage 4 were used for the screen. Prior to seeding, cells were detached using Trypsin-EDTA (Gibco, 0.05%), counted using a hemocytometer, and seeded at 1,053 cells/cm² (2,000 cells per well) onto topography-containing 24-well plates. After seeding, the plates were allowed to rest undisturbed in the biosafety cabinet for 10 minutes to facilitate uniform cell attachment before transfer to the incubator. Cells were cultured for 48 hours without medium change prior to fixation. Ten unique surface designs were used with one surface per design for four surfaces, and two technical replicas for 6 surfaces. All steps were conducted under sterile conditions and were detailed in a publicly available protocol. (https://github.com/cbite/Micellangelo/tree/main/Raw_data_and_preprocessing/Wet_lab_work_protocols)

### 2.4. Fixation, immunostaining and mounting

Following 48 hours of culture, cells were washed three times with PBS (1 mL) and fixed in-well using paraformaldehyde (PFA; Thermo Fisher, methanol-free, 0.5 mL 4%) for 20 minutes at room temperature and washed three times with sterile PBS. Permeabilization was performed using Triton X-100 (Merck, 0.5%) in PBS for 15 minutes, followed by 3 times washing with PBS and blocking in blocking buffer (bovine serum albumin, Roche Diagnostics GmbH, 3%) and glycine (VWR Chemicals, BDH Chemicals, 0.3 m) in PBS) for 45 minutes at room temperature. Primary staining was performed right after blocking by incubating cells overnight at 4°C with mouse anti-YAP monoclonal antibody (Santa Cruz, sc-101199, lot #G3119) diluted in blocking buffer (1:500). The following day, cells were washed three times with PBS and incubated for 1 hour at room temperature with Alexa Fluor 488-conjugated goat anti-mouse IgG2a secondary antibody (Invitrogen, A21131, 1:400) diluted in PBS. Actin cytoskeleton was labelled using phalloidin (Abcam, ab176756 or ab176759) diluted in PBS (1:1000) for 15 minutes. Nuclei were counterstained with DAPI (Invitrogen, D1306; 1:1000) diluted in PBS for 8 minutes. All staining steps were separated by three PBS washing steps. A drop of Mowiol 4-88 mounting medium (15 μL) was gently pipetted directly onto clean (wiped with 70% ethanol, 15 minutes drying) glass slides. (Microscope Slides, Cut Frosted, Epredia New Erie Scientific LLC, Cat. No. AA00000112E01MNZ10, Lot 06322, 76×26×1.0 mm). Topographic inserts were carefully retrieved from the well using fine tweezers and placed on the Mowiol droplet, cell side facing up. Another Mowiol drop (15 μL) was added onto the surface, and a glass coverslip (Cover Slips, 24 × 60 mm, No. 1.5, Epredia, Epredia Netherlands B.V.) was then lowered onto the sample, starting from one edge to avoid air bubbles. After positioning, the corners of the coverslip were sealed using clear nail polish (Etos b.v.) to prevent dehydration and maintain mounting stability. Samples were stored at 4°C, protected from light, and imaged between 30 and 100 days post-staining.

### 2.5. Microscopy and image acquisition

Fluorescence and brightfield imaging were performed on a Nikon Ti2-E inverted high-throughput screening (HTS) microscope equipped with a CFI S Plan Fluor ELWD 20×C air objective (NA 0.45, WD 8.1-6.9 mm). The microscope was controlled via Nikon NIS-Elements software, and all image acquisition was performed using a motorized stage and automated large-field scanning.

Prior to high-resolution imaging, each surface was pre-screened at 10× magnification to assess staining quality, signal-to-noise ratio, and the presence of sufficient numbers of non-touching single cells. Staining artifacts were defined as regions with high background fluorescence or low target signal. Only samples with uniform staining across DAPI, phalloidin, and YAP channels and low fluorescent background were selected for 20× imaging. Exposure settings for each fluorescence channel were adjusted using the NIS-Elements “Highlight Overexposed Pixels” plugin to limit saturation to less than 1% of image pixels. This standard was uniformly applied across all samples to preserve fluorescence intensity for downstream quantification **(Supplementary Table 1)**.

To account for variations in sample tilt, an autofocus heatmap was generated for each surface using the phalloidin fluorescence signal. Autofocus anchor points were established by manually setting focus at four equidistant points across the surface edge, as well as at the center of the surface, to ensure consistent focal planes across all topographies. Regions of interest (ROIs), the coordinates defining the area to be imaged, were defined using the 3-Point Rectangle Tool, ensuring complete coverage of the imaged surfaces while maintaining a consistent 0.5 millimeter margin around each insert.

Phalloidin staining for actin was imaged using either TRITC or Cy5 channels depending on fluorophore availability. Fluorescence and brightfield channels were acquired sequentially using the following configuration:

**Supplementary Table 1.** Fluorescence imaging channels, fluorophores, and filter settings used in high-throughput microscopy.

| Channel | Target | Fluorophore | Excitation (nm) | Emission (nm) |
| --- | --- | --- | --- | --- |
| c01 | Actin | TRITC (Phalloidin 555) or Cy5 (Phalloidin 647) | 540–568 or 590–645 | 579–640 or 659–736 |
| c02 | YAP | FITC (Alexa Fluor 488) | 461–488 | 503–548 |
| c03 | Nuclei | DAPI | 379–405 | 414–480 |
| c04 | Brightfield | DIA-PH3 | — | — |

Large-field acquisition was performed with 10% image overlap and 1×1 binning to preserve spatial resolution. Stitching was handled automatically by the NIS-Elements large image acquisition module. Image files were saved as .nd2 format with a standardized filename structure [(User_ID)_FeatureIdx(X)_YAP_PHA_DAPI.nd2]. A summary of the cell culture, fixation, and imaging metadata, including details on surface topography identifiers, emission wavelengths, and extracted single-cell counts, is provided in Supplementary Table 2 below.

**Supplementary Table 2.**
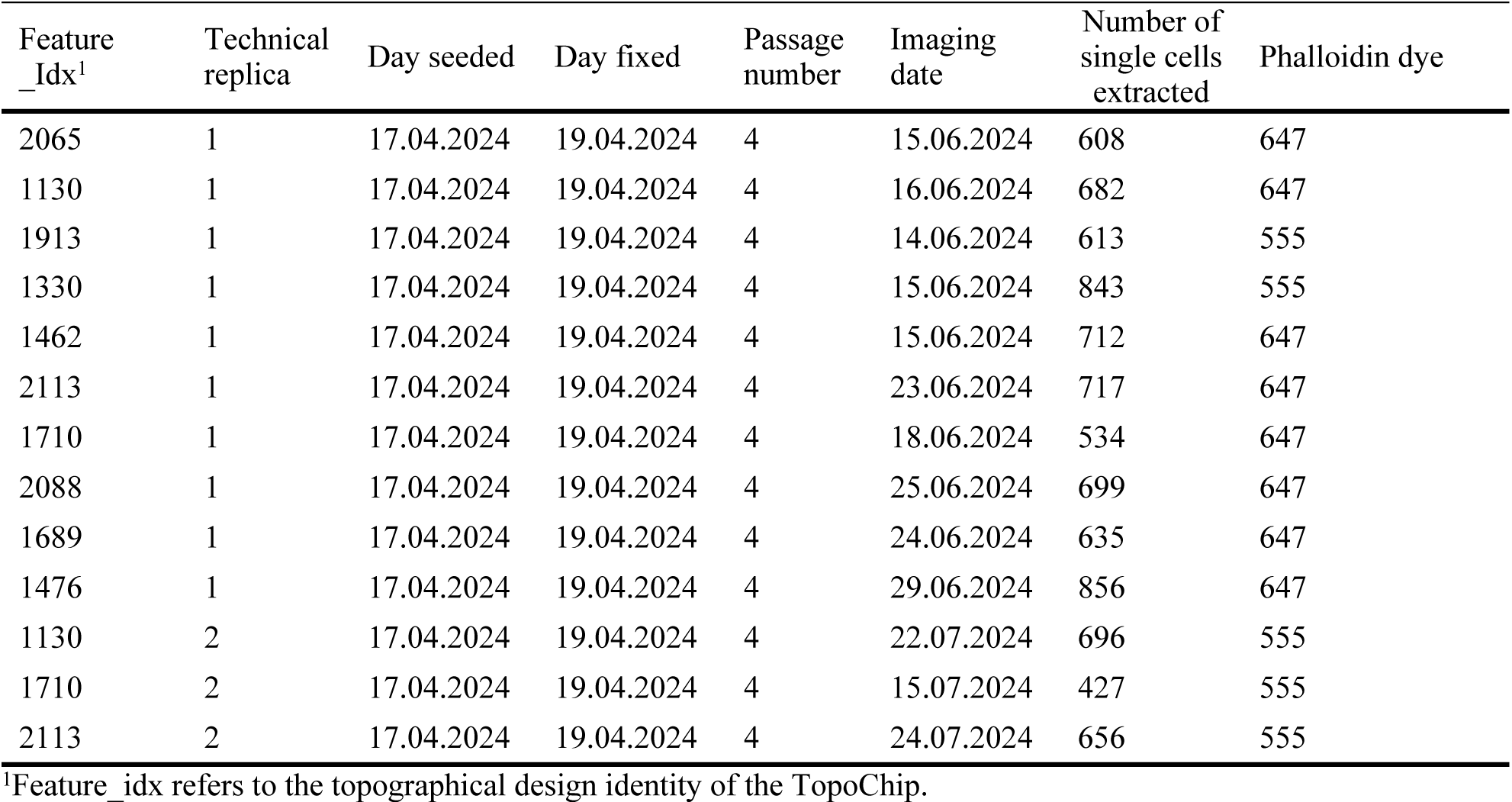
Summary of cell culture, fixation, and imaging metadata for individual topographies.

| Feature<br>_Idx<sup>1</sup> | Technical<br>replica | Day seeded | Day fixed | Passage<br>number | Imaging<br>date | Number of<br>single cells<br>extracted | Phalloidin dye |
| --- | --- | --- | --- | --- | --- | --- | --- |
| 2065 | 1 | 17.04.2024 | 19.04.2024 | 4 | 15.06.2024 | 608 | 647 |
| 1130 | 1 | 17.04.2024 | 19.04.2024 | 4 | 16.06.2024 | 682 | 647 |
| 1913 | 1 | 17.04.2024 | 19.04.2024 | 4 | 14.06.2024 | 613 | 555 |
| 1330 | 1 | 17.04.2024 | 19.04.2024 | 4 | 15.06.2024 | 843 | 555 |
| 1462 | 1 | 17.04.2024 | 19.04.2024 | 4 | 15.06.2024 | 712 | 647 |
| 2113 | 1 | 17.04.2024 | 19.04.2024 | 4 | 23.06.2024 | 717 | 647 |
| 1710 | 1 | 17.04.2024 | 19.04.2024 | 4 | 18.06.2024 | 534 | 647 |
| 2088 | 1 | 17.04.2024 | 19.04.2024 | 4 | 25.06.2024 | 699 | 647 |
| 1689 | 1 | 17.04.2024 | 19.04.2024 | 4 | 24.06.2024 | 635 | 647 |
| 1476 | 1 | 17.04.2024 | 19.04.2024 | 4 | 29.06.2024 | 856 | 647 |
| 1130 | 2 | 17.04.2024 | 19.04.2024 | 4 | 22.07.2024 | 696 | 555 |
| 1710 | 2 | 17.04.2024 | 19.04.2024 | 4 | 15.07.2024 | 427 | 555 |
| 2113 | 2 | 17.04.2024 | 19.04.2024 | 4 | 24.07.2024 | 656 | 555 |

<sup>1</sup>Feature\_idx refers to the topographical design identity of the TopoChip.

### 2.6. Image preprocessing & conversion

All image stacks acquired on the Nikon Ti2-E microscope were initially saved in .nd2 format to preserve full metadata and multi-channel fluorescence information. To enable downstream analysis in open-source environments, .nd2 files were converted into multi-channel .tif format using the *Nikon ND2 Reader* plugin in Fiji (ImageJ, version 1.54f). Conversion was performed without any form of contrast adjustment, gamma correction, or fluorescence signal enhancement. The exported filenames adhered to the following structure:

Channel[Fluor]-[Index]_DIA-[PhaseIndex],DAPI2_Seq[SequenceNumber]_[Fluor].tif

This naming scheme encoded the fluorescence channel (e.g., TRITC, FITC), phase contrast references (e.g., DIA-Ph3), sequence ID (not used in these experiments) and the specific stain (e.g., DAPI2), providing robust traceability for downstream channel-specific processing and QC. All .tif files were stored alongside their corresponding raw .nd2 files on a local research server. Files were organized into structured directories sorted by date, and featureidx. Folder-level metadata included information on surface identity, replicate ID, and imaging dates (Supplementary Table 2).

### 2.7. Image tiling and field selection

Stitched .tif images were tiled into 2000×2000 pixel-sized tiles using a custom MATLAB (R2022a) script, resulting in around 120 images per topography insert. Next, tile quality was visually assessed and exclusion criteria were: tiles outside or spanning the border of the topographical area and images exhibiting obvious imaging brightfield artifacts. In the process, the observed topographical pattern was verified against the design file. A subset of high-quality brightfield tiles was selected base on absence of manufacturing artifacts, absence of dust particles, air bubbles and on uniform illumination.

### 2.8. CellProfiler feature extraction

Following tile extraction, segmentation of individual cells was performed using a pipeline developed in CellProfiler version 4.2.5, ^[65]^ which performed robustly across most chips. For certain chips, this pipeline was manually adjusted to account for surface-specific variation. Each resulting pipeline version was saved individually (https://github.com/cbite/Micellangelo/blob/main/Raw_data_and_preprocessing/cellprofiler_p rotocols_for_all_the_chips.zip). Conceptually, the pipeline worked as follows:

1. Illumination correction – applied individually to each image to correct for uneven background lighting and staining artifacts, such as vignetting or intensity gradients.
2. Object segmentation with nuclei (DAPI) as primary object and cell body (phalloidin) as the secondary object.
3. Removal of all touching objects as well as objects crossing tile borders.
4. Measurements (location, intensity distribution and morphological features).
5. Data exportation.

Before execution, for each surface, intensity thresholds were assessed using Test mode to ensure robust object segmentation. Prior to execution we ensured that all necessary output objects were selected for export. During the export stage the ExportToSpreadsheet module was used, that allows downstream quality control with *CellProfiler Analyst*. ^[66]^ There, for location referencing the non-touching nuclei objects were chosen.

Each customized pipeline was saved under the directory: cellprofiler_protocols_for_all_the_chips/[CHIPNUMBER]/ with filenames including the chip identifier in the format: Feature_Idx_[CHIPNAME].cpproj

Upon successful execution, the output measurements were saved following a structured hierarchy in the main data repository: …/[CHIPNAME]/CP output/ – for all exported measurements, masks, and database files.

### 2.9. Segmentation quality control

For each surface, a CellProfiler Analyst based classifier was trained to classify objects as well segmented or badly segmented. 30 objects were randomly sampled and classified as either good or bad. Next, about 100 good objects were retrieved from the database, and visually classified as good/bad, and the same was performed for negative objects. Finally, the classifier was asked to retrieve every uncertain object, which were then also manually classified as good/bad. Then, the classification algorithm was considered trained, and all objects were assigned their class. The resulting classification tables were exported as: QC_classifier_table.csv. Negative objects (those marked as failed by the classifier) were programmatically excluded during concatenation. Final filtering was based on cellular morphology metrics. For each cell, the logarithm of nuclear-to-cytoplasmic area ratio (NtoC) was calculated by dividing the segmented area of the nucleus (from the DAPI channel) by that of the cytoplasm (from the phalloidin channel). This ratio served as a sensitive indicator of segmentation fidelity and biological plausibility.^[67]^ Outlier cells were identified using an interquartile range (IQR) filter: cells with NtoC log-area ratios falling outside the IQR ± 1.5× range were flagged for potential exclusion. Each flagged object was visually inspected to ensure that only genuinely mis-segmented or biologically implausible cells were removed, while atypical but valid morphologies were retained. The result was a refined dataset composed of high-confidence, well-segmented single fibroblasts suitable for downstream morphometric and phenotypic analysis. Supplementary Table 3 summarizes the results of the quality control procedure.

**Supplementary Table 3.**
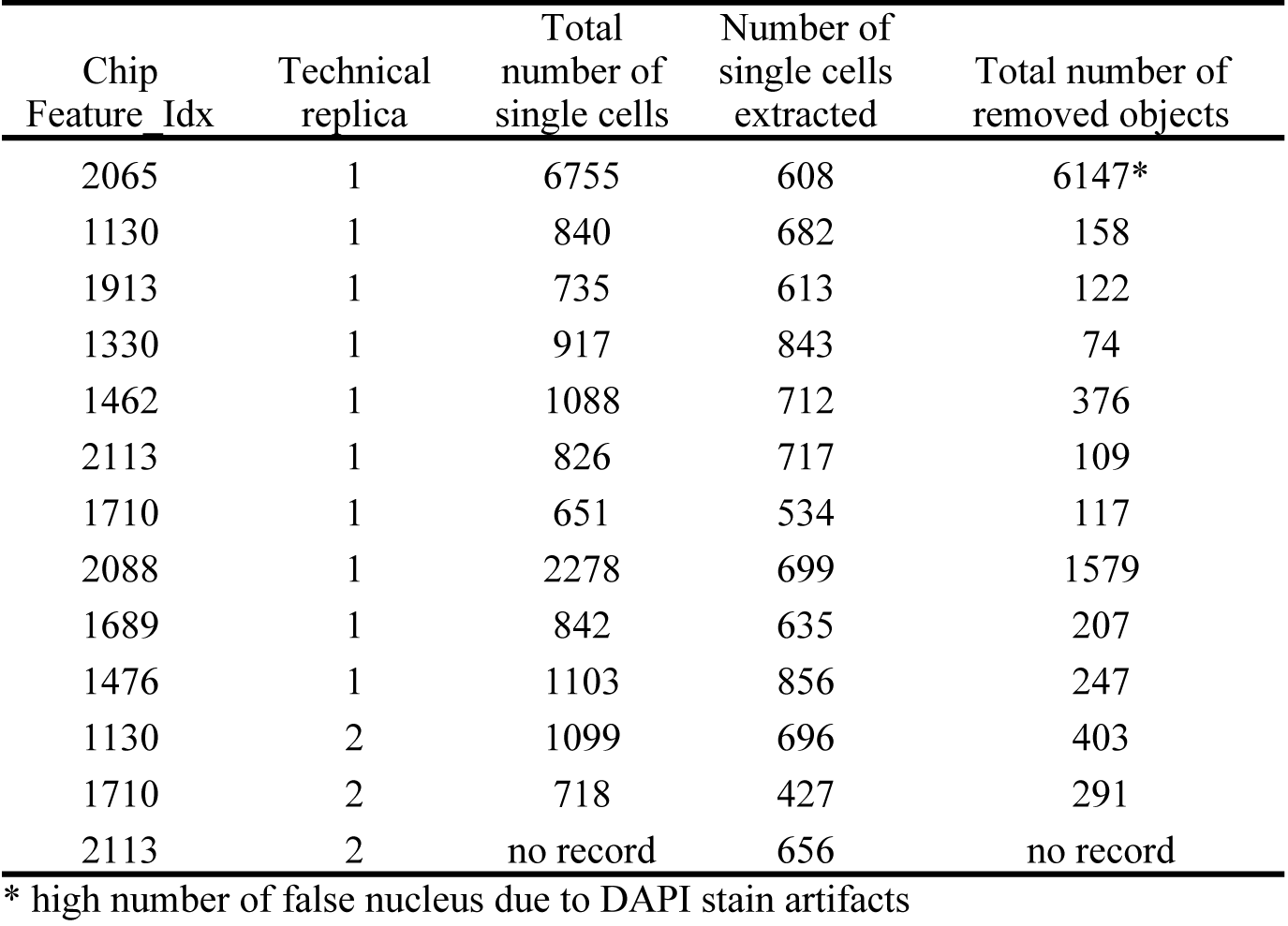
Per-Chip Summary of Single-Cell Segmentation QC, Classification, and Filtering Outcomes.

| Chip<br>Feature Idx | Technical<br>replica | Total<br>number of<br>single cells | Number of<br>single cells<br>extracted | Total number of<br>removed objects |
| --- | --- | --- | --- | --- |
| 2065 | 1 | 6755 | 608 | 6147* |
| 1130 | 1 | 840 | 682 | 158 |
| 1913 | 1 | 735 | 613 | 122 |
| 1330 | 1 | 917 | 843 | 74 |
| 1462 | 1 | 1088 | 712 | 376 |
| 2113 | 1 | 826 | 717 | 109 |
| 1710 | 1 | 651 | 534 | 117 |
| 2088 | 1 | 2278 | 699 | 1579 |
| 1689 | 1 | 842 | 635 | 207 |
| 1476 | 1 | 1103 | 856 | 247 |
| 1130 | 2 | 1099 | 696 | 403 |
| 1710 | 2 | 718 | 427 | 291 |
| 2113 | 2 | no record | 656 | no record |

\* high number of false nucleus due to DAPI stain artifacts

### 2.10. Standardized coordinate system

Due to the spatially repeating nature of the surface topography, cells positioned at different absolute locations within an image but the same relative offset to the topographical motif experienced near-identical environments. To eliminate redundant variation in equivalent spatial contexts, a canonical coordinate system was defined per insert. For each surface, a 750×750 pixel snippet of the brightfield image, free from production or imaging artifacts, was selected to serve as the reference frame. Following segmentation quality control, we aligned the brightfield channel of each single-cell image to this standardized reference using phase cross-correlation, implemented via the skimage.registration.phase_cross_correlation function (https://scikit-image.org/docs/0.25.x/api/skimage.registration.html#skimage.registration.phase_cross_correlation) in scikit-image (v0.25). This alignment process enabled consistent positioning of cells relative to the topographical features. Visual inspection of the registered outputs confirmed that the majority of cells were centrally located within the coordinate system, while maintaining correct relative alignment to the background topography.

